# Signal Peptide of HIV-1 Envelope Modulates Glycosylation Impacting Exposure of V1V2 Epitopes

**DOI:** 10.1101/2020.07.21.212183

**Authors:** Chitra Upadhyay, Roya Feyznezhad, Liwei Cao, Kun-Wei Chan, Kevin Liu, Weiming Yang, Hui Zhang, Jason Yolitz, James Arthos, Arthur Nadas, Xiang-Peng Kong, Susan Zolla-Pazner, Catarina E. Hioe

**Affiliations:** Icahn School of Medicine at Mount Sinai, New York, NY, USA; James J. Peters Veterans Affairs Medical Center, Research Service, Bronx, NY, USA; Johns Hopkins University, Baltimore, MD USA; Department of Biochemistry and Molecular Pharmacology New York University School of Medicine, New York, USA; Laboratory of Immunoregulation, National Institute of Allergy and Infectious Diseases, National Institutes of Health, Bethesda, MD USA; Institute of Environmental Medicine, New York University School of Medicine, New York, New York, USA

**Keywords:** HIV, HIV envelope, signal peptide, glycosylation, neutralization, V1V2 antibodies, trimer apex

## Abstract

HIV-1 envelope (Env) is a trimer of gp120-gp41 heterodimers, synthesized from a precursor gp160 that contains an ER-targeting signal peptide (SP) at its amino-terminus. Each trimer is swathed by ∼90 N-linked glycans, comprising complex-type and oligomannose-type glycans, which play an important role in determining virus sensitivity to neutralizing antibodies. We previously examined the effects of single point SP mutations on Env properties and functions. Here, we aimed to understand the impact of the SP diversity on glycosylation of virus-derived Env and virus neutralization by swapping SPs. Analyses of site-specific glycans revealed that SP swapping altered Env glycan content and occupancy on multiple N-linked glycosites, including the conserved N156 and N160 glycans in the V1V2 region at the Env trimer apex. Virus neutralization was also affected, especially by antibodies against the V2i, V2p and V2q epitopes. Likewise, SP swaps affected the recognition of soluble and cell-associated Env by antibodies targeting distinct V1V2 configurations. These data highlight the contribution of SP sequence diversity in shaping the Env glycan content and its impact on the configuration and accessibility of V1V2 epitopes on Env.

**Author Summary:** HIV-1 Env glycoprotein is produced by a precursor gp160 that has a signal peptide at its N-terminus. The SP is highly diverse among the HIV-1 isolates and no two SP are same. This study presents site-specific analyses of N-linked glycosylation on HIV-1 envelope glycoproteins from infectious viruses produced with different envelope signal peptides. We show that signal peptide swapping alters the envelope glycan shield, including the conserved N156 and N160 located in the V1V2 region on the trimer apex, to impact Env recognition and virus neutralization by antibodies, particularly those targeting the the V1V2 region. The data offer crucial insights into the role of signal peptide in the interplay between HIV-1 and antibodies and its potential utility to control Env glycosylation in the development of Env-based HIV-1 vaccine.

## Introduction

The HIV-1 envelope glycoprotein (Env) spike is the only viral protein accessible to neutralizing antibodies; therefore, a critical HIV-1 vaccine component. Env is synthesized as a precursor gp160 glycoprotein, which is targeted to the endoplasmic reticulum (ER) by the signal peptide (SP, also known as signal or leader sequence) present at the amino terminus of the nascent protein. Generally SPs contain ∼16-30 amino acids with a characteristic tripartite structure: a hydrophilic positively charged n-region, a central hydrophobic h-region, and a slightly polar carboxy terminal c-region with a cleavage site for signal peptidase (1). SPs are typically cleaved from the maturing protein co-translationally, but the ∼30 amino-acid long SP of HIV-1 Env is cleaved post-translationally (2-5). This delayed SP cleavage is conserved across HIV-1 subtypes and functions as a quality-control checkpoint, ensuring proper folding and post-translational modifications of gp160 before its egress from the ER (6).

Within the ER, gp160 is decorated with ∼30 N-linked glycans and undergoes folding and extensive isomerization until near native conformation is reached with the SP still attached. The SP is cleaved before gp160 reaches the Golgi for further glycosylation maturation and other post-translational modifications (2, 4, 7). In the Golgi, gp160 is cleaved by cellular furin to generate non-covalently linked gp120-gp41 heterodimers, three of which assemble to form the functional Env spikes. The apex of the trimer is made of three V1V2 domains from the gp120 protomers with three V3 loops laying underneath and partially concealed in the native “closed” conformation (8-10). V1V2 epitopes are of particular interest, since the RV144 Thai vaccine trial identified high levels of V1V2-specific antibodies as a correlate of reduced infection risk (11-13). While V1V2 and V3 contain cross-reactive immunogenic epitopes, these regions are structurally dynamic and the epitopes are conformationally masked in most HIV-1 isolates (14). Dense glycosylation further shields V1V2, V3 and other epitopes (15-17). Nonetheless, some glycans also serve as components of epitopes for broadly neutralizing antibodies (bnAbs) (18-21).

Each V1V2 domain can form a 5-strand beta-barrel module, with the polymorphic V2 C-strand adopting α-helical or β-sheet conformations (22). Three types of V1V2 epitopes have been defined: V2p, V2i and V2q (23). The V2p type constitutes the α-helical peptide epitopes in the V2 C-strand that are recognized by monoclonal antibodies (mAbs) CH58 and CH59 from RV144 vaccinees (24) and the CAP228 mAb series from a virus-infected donor in South Africa (25). The V2i-type epitopes are made of conformational discontinuous segments in V1V2, some of which are near the integrin α4β7-binding motif (26, 27); these epitopes are recognized by mAbs such as 830A, 697-30D, and 2158 from infected US patients (26). The V2q type is comprised of quaternary neutralizing epitopes targeted by bNAbs, including the PG9 and PG16 (28). The V2q epitopes requires that the V2 C-strand assume a beta-sheet structure (19). The V2q epitopes also include glycans at positions N156 or N173 and N160 on the V1V2 apex (19, 29), which, with N197 and N276 glycans creates the conserved ‘trimer-associated mannose patch’ (TAMP). Other Abs such as mannose-binding 2G12 and some PGT bnAbs (e.g. PGT121, PGT128, PGT135) also directly target glycans at N295, N332, and N392 (30-32), which form the intrinsic mannose patch (IMP) (33). The V2i epitopes, in contrast, do not include glycans, but are conformationally dependent on proper V1V2 glycosylation and folding, and require the presence in the V2 C-strand of a beta-strand configuration (22, 23). Moreover, the propensity of V2 C-strand to adopt α-helical and β-sheet structures recognized by V2p and V2q mAbs, respectively, is influenced by glycosylation (23). Thus, changes in glycosylation can impact HIV-1 interaction with Abs against V1V2. This would be expected to be true for other Env regions as well.

We and others have recently shown that HIV-1 Env glycosylation is dictated in part by the SP (34, 35). The SP displays remarkable sequence diversity (34, 36, 37), the reason for which are not fully understood. Nonetheless, the native Env SP has been replaced by unrelated SPs (e.g. SPs of CD5 and tissue plasminogen activator (tPA)) to promote the secretion of soluble Env (5, 37-39). Likewise, the herpes simplex virus (HSV) SP was used to express the gp120 tested in RV144 clinical trial (40-42); the only trial with a modest 31 % protective efficacy (40-42). Beyond protein targeting, SP also influences protein folding, modification, localization, and topology (3, 43-45). Mutations in SPs are suggested to impair transport and alter maturation as a result of incomplete glycosylation in the ER and Golgi, leading to several human diseases (46). For example, a mutation in the hydrophobic core of the SP of preproparathyroid hormone is suggested to impact its translocation across the ER to cause a functional defect associated with familial isolated hypoparathyroidism (46). Mutations in the SP of TGF-beta type I receptor results in resistance to TGF-beta in B cell chronic lymphocytic leukemia patient (47). SP mutations also have been associated with other metabolic, developmental, cardiovascular, and autoimmune human diseases (47-51). The effect of SP and its residues on viruses has not been much studied. The extremely long Env SPs of retroviruses like foamy virus and mouse mammary tumor virus have been shown to mediate additional functions other than ER targeting. The cleaved SP product from foamy virus is associated with cell-free virions and is critical for virus release from infected cells (52). The SP of the MMTV envelope precursor is reported to function as a nuclear export factor for intron-containing transcripts (53). Expressing HIV-1 gp120 with SPs from heterologous isolates also altered gp120 glycosylation and reactivity with lectins and mAbs (35). Collectively, these studies highlight the significant role of SPs and their profound effects on the functions of associated proteins and viruses.

While the significance of sequence diversity in the HIV-1 gp120 and gp41 subunits on antibody recognition has been extensively documented, much less information is available concerning the sequence variations in the HIV-1 Env SP. We previously reported that the HIV-1 Env glycan content can be altered by single point mutations in the Env SP to affect virus sensitivity to neutralization by antibodies, although only two out of 29 potential N-linked glycosites (PNGSs) were detected (34). In this study we identified an array of PNGSs impacted by exchanging the entire native Env SP with SPs from heterologous HIV-1 isolates in the proviral context. Using Env backbones and SPs from viruses representing different clades, neutralization tiers and clinical stages (S1 Fig), we tested the impact of SP exchanges on Env functions in the setting of different Env backbones. Site-specific glycosylation analysis of virion-derived Env expressed with SPs from native versus heterologous HIV-1 strains revealed altered proportions of oligomannose and complex glycans at many glycosites including the highly conserved N156 and N160 glycans on the V1V2 trimer apex. The modified glycan compositions were associated with perturbed virus neutralization and Env binding, especially by antibodies targeting V1V2. Because SP swapping is a common strategy employed to construct Env-based HIV-1 vaccines, this study provides, previously unknown, details about the critical contribution of SP in affecting HIV-1 Env properties with important implications for vaccine designs.

## Results

### Glycosylation changes of SP-swapped vs WT Envs on CMU06 virions

To investigate the functional consequences of Env SP diversity, we selected SPs from isolates that display different neutralization sensitivity and represent different clades and clinical stages (37, 54, 55). The SP of CMU06 Env (CRF01_AE, tier 2, acute) was swapped with SPs from MW (clade C, tier 1A, chronic), 398F1 (clade A, tier 1A, acute), CH119 (CRF_07, tier 2, chronic) and 271.1 (clade C, tier 2, acute), while maintaining the SP cleavage site of the parental CMU06 (Fig 1A). All SP-swapped viruses were infectious and had comparable levels of Env incorporated into virions (S2A and S2B Fig), demonstrating that SP swapping had no deleterious effect on Env expression and virus infectivity.

**Figure 1.**
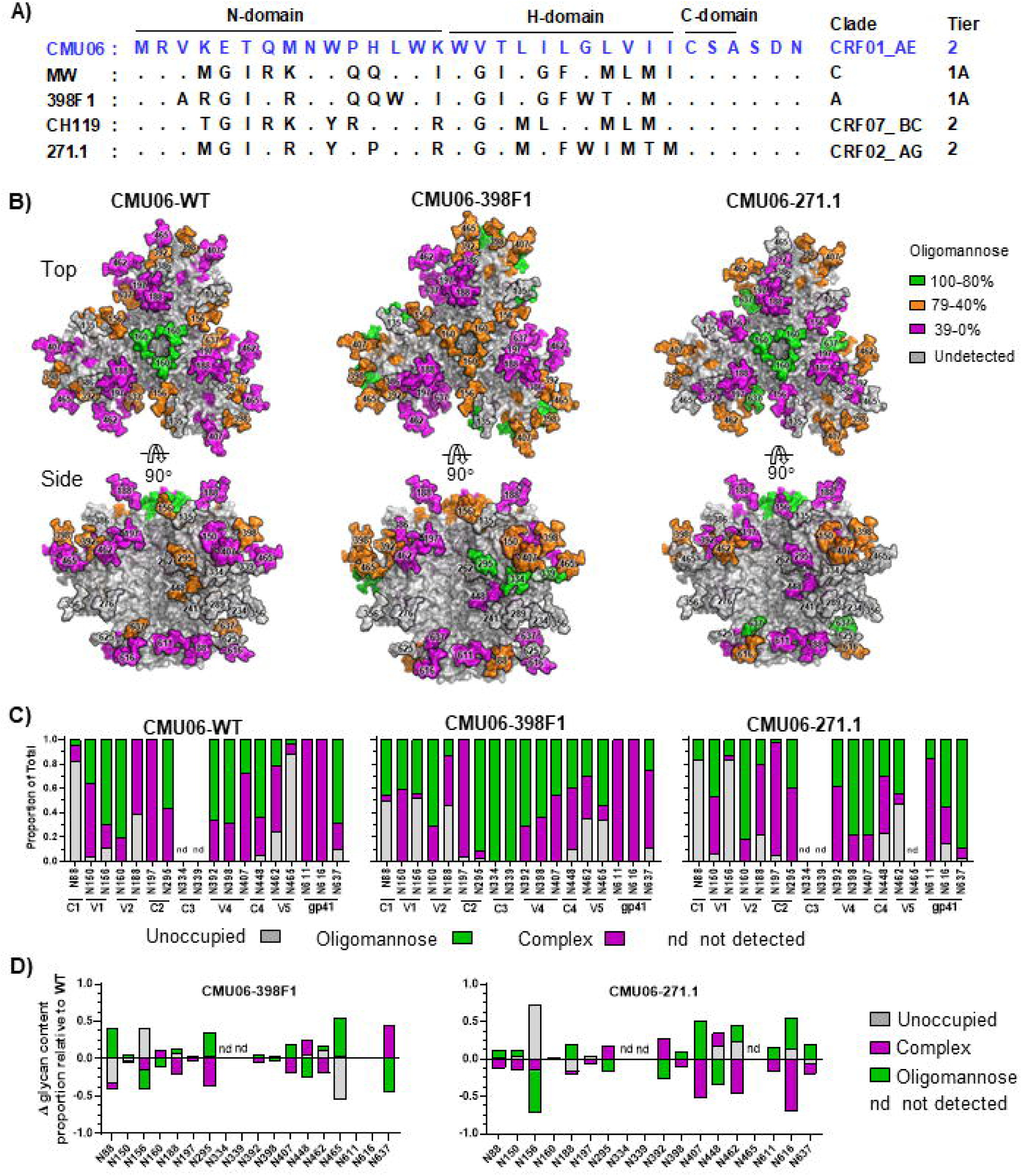
Effect of SP swap on CMU06 Env glycosylation. (A) CMU06 WT and swapped SPs. SP cleavage site is denoted by red triangle. Within the alignment, dots indicate identity with the WT SP sequence. (B) Env trimer models (generated using a SOSIP.664 trimer PDB 5FYJ) showing the predominating glycoforms at each of the CMU06 glycosites for CMU06-WT, CMU06-398F1, and CMU06-271.1. Type of glycans added was based on LC-MS/MS data of virion-associated Envs from 293T cells (Fig 1C). (C) Relative amounts of unoccupied, oligomannose, and complex glycans at glycosites on SP-swapped vs WT CMU06 as determined by LC-MS/MS. nd: glycosites not detected in 1 or 2 Envs. (D) Changes in glycan content at each glycosite for CMU06-398F1 and CMU06-271.1 vs CMU06-WT calculated from Fig 1C data.

We initially appraised the impact of SP swapping on glycosylation by subjecting the Envs from sucrose-pelleted WT vs SP-swapped CMU06 viruses to Western blots probed with anti-gp120 mAbs and lectins that recognize distinct sugar moieties: GNA (terminal α1-3 and α1-6 mannose on high-mannose glycans) and AAL (α1-6 or α1-3 fucose on complex glycans) (56, 57). Upper and lower Env bands were detected by anti-gp120 mAbs and lectins (S2C Fig). GNA binding to the upper bands of SP-swapped Envs was increased by ∼2-fold vs WT, whereas binding to the lower bands were minimally altered (S2C Fig). Differences were also observed in AAL binding to the upper and lower bands of all SP-swapped Envs. These data demonstrated modulations in glycan contents of SP-swapped Envs vs WT that were modest but enriched for terminal α1-3 and α1-6 mannose-bearing oligomannose-type glycans and fucosylated complex glycans.

To further discern glycan alterations at specific N-linked glycosites, CMU06-WT, CMU06-398F1, and CMU06-271.1 were analyzed by LC-MS/MS on an Orbitrap MS1 spectrometer upon Endo-H and PNGase digestion. Of 28 potential glycosites present on CMU06 Env (S3 Fig), oligomannose glycans, complex glycans, and unoccupied glycosites were identified with coverage of 57% (16/28), 64% (18/28), and 54% (15/28) respectively. Hybrid-type glycans were indistinguishable from oligomannose-type, as both were digested by Endo-H. Noticeable alterations were found at many glycosites in both gp120 and gp41 of CMU06-398F1 and CMU06-271.1 vs CMU06-WT, (Fig 1B-E). Of the 12 glycosites detected on gp120 from all 3 Envs, the percentage of oligomannose glycans was same on CMU06-WT (41%) and CMU06-271.1 (41%) but changed to 47% on CMU06-398F1. Of the three gp41 glycans, complex types were predominant on WT (74%) and the amounts were increased to 88% on 398F1 and decreased to 41% on 271.1. Consistent with GNA binding data (S2C Fig), LC-MS/MS data demonstrated an increase in total oligomannose glycans of CMU06-398F1 (40%) and CMU06-271.1 (43%) vs CMU06-WT (35%). The data also showed alterations in the total complex glycans (45%: 398F1 and 38%: 271.1 vs 48%: WT), but LC-MS/MS could not delineate the specific proportion of fucose-bearing glycans detected by AAL.

A model of glycosylated trimeric Env was generated to illustrate glycosite-specific alterations on CMU06-398F1 and CMU06-271.1 vs CMU06-WT (Fig 1B-D). Prominent changes were observed in glycan contents of N156 and N160 at the trimer apex (Fig 1B-E). N156 and N160 glycans constitute the TAMP and serve as key contact residues for bnAbs such as PG9, PG16, PGT145 and PGDM1400 (19, 58, 59). On CMU06-WT Env, N156 was occupied by oligomannose (70%) and complex (19%) glycans, while 11% was unoccupied. In contrast, N156 was largely unoccupied on CMU06-398F1 (52%) and CMU06-271.1 (83%). When N156 of CMU06-398F1 was occupied (48%), oligomannose glycans predominated at 45% and complex glycans constituted only 3%. The N173 glycan, which is recognized in place of N156 by PG9 on some HIV-1 isolates (60), is absent on CMU06 (S3 Fig). N160 was decorated mostly by oligomannose glycans in CMU06-WT and CMU06-271.1 (81-82%) but had reduced abundance of oligomannose in CMU06-386F1 (71%). Complex-type glycans increased proportionally to 29% at N160 of CMU06-386F1 vs 19% in CMU06-WT and 18% in CMU06-271.1. At the nearby N188, also emanating from the apical V1V2 region, 398F1 and 271.1 SP swapping increased oligomannose glycans to 13% and 20%, respectively, from 0% in CMU06-WT.

Additional changes were also noted in glycans projecting laterally from the Env core at positions N295, N392, N407, N448, N462, and N465. N295 and N392, which together with N332 or N334 comprise the IMP, had lower or higher proportions of complex glycans in SP-swapped viruses vs WT, whereas N334 was detected in only CMU06-398F1 to contain oligomannose glycans. However, not all glycosites were affected by SP swapping; minimal to no changes were observed at N197, 94-100% of which was comprised of complex glycans, and at N398, which contained complex and oligomannose glycans at a ratio of ∼1:2 in all three Envs tested.

At the trimer base, N88 was largely unoccupied in CMU06-WT (82%) and CMU06-271.1 (82%) but had 50% occupancy in CMU06-398F1 with mainly oligomannose glycans. N611 and N616 at the trimer base in gp41 were 100% occupied by complex glycans in CMU06-WT and CMU06-398F1, but in CMU-271.1 oligomannose glycans were found at N611 (16%) and N616 (55%). The oligomannose content of N637 in gp41 increased from 69% in CMU06-WT to 89% in CMU06-271.1 but was reduced to 25% in CMU06-398F1. Glycosites at N135, N234, N241, N262, N276, N289, N356, N386 and N625 were undetectable in all three Envs. Collectively, the data provides detailed insight into the role of SP in influencing Env glycan composition.

### Altered mAb recognition of CMU06 as a result of SP swapping

Next, we examined the effect of SP swap on virus neutralization. Changes in sensitivity to neutralization were observed with each SP-swapped virus vs WT especially by V1V2 mAbs (Fig 2A-B). CMU06-CH119 and CMU06-271.1 were more resistant to neutralization by most V2i mAbs tested. Increased sensitivity to V2q mAbs PG9 and/or PG16 was observed for CMU06 with SPs from MW, 398F1, and 271.1. Neutralization of CMU06-WT by V3 crown-specific mAbs 2219 and 2557 was weak (IC50 >100 ug/ml), but switching SPs further reduced neutralization by 2219 and 2557. Neutralization by V2p mAb (CH59), gp41-MPER mAb (2F5), CD4bs mAbs, and CD4-IgG was not much affected (Fig 2A-B).

**Figure 2.**
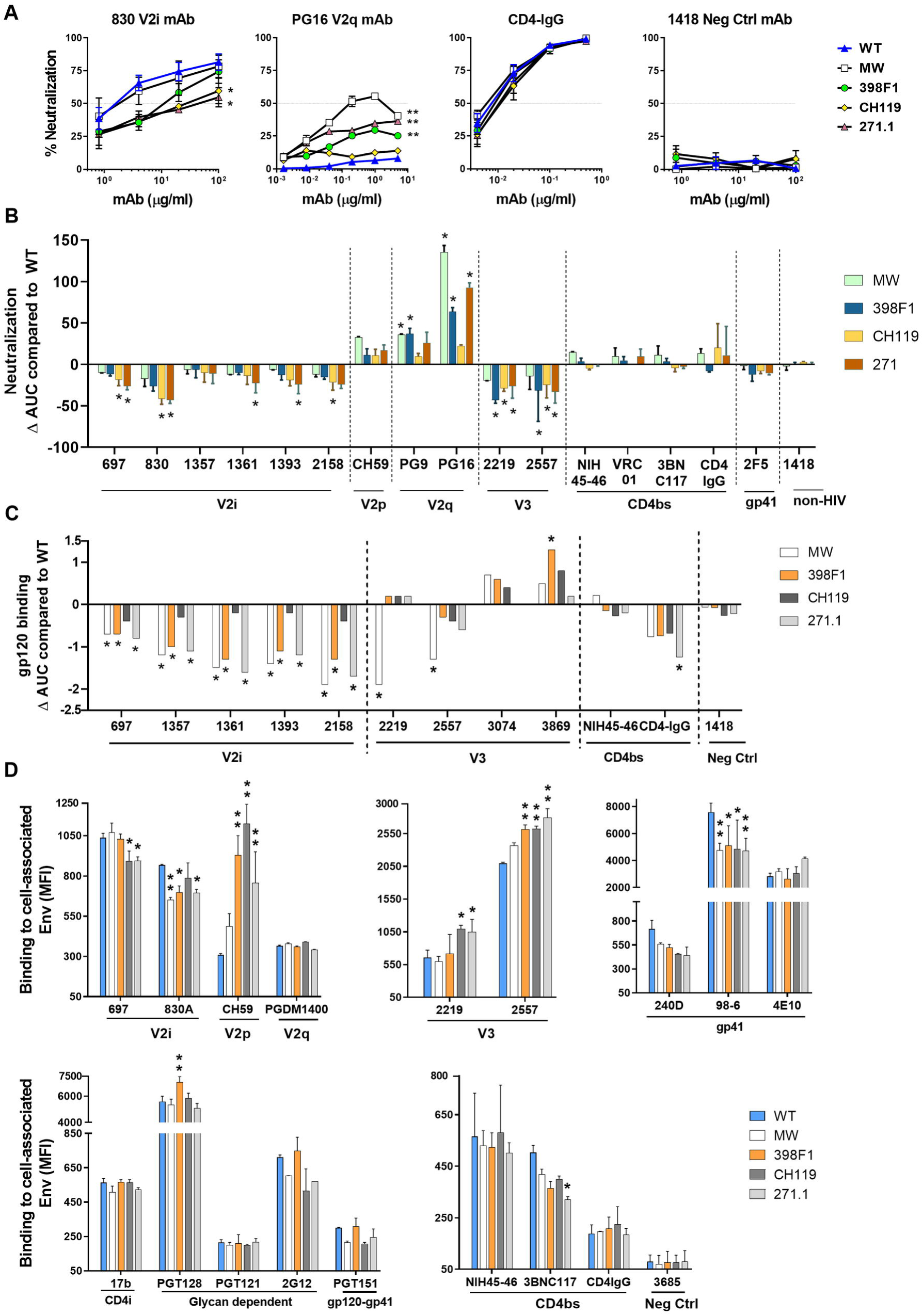
Effect of SP swap on CMU06 Env-antibody interaction. (A) Neutralization of 293T-derived SP-swapped vs WT CMU06. Viruses were incubated with titrated mAbs at 37°C for 24 hours before addition of TZM.bl cells, except for PG9, NIH45-46, VRC01, 3BNC117, 2F5 and CD4-IgG which were pre-incubated with viruses for 1 hour. MAb 1418 served as negative control. Representative neutralization curves are shown. Mean and SEM from 2-3 experiments are shown. Data were analyzed by 2-way ANOVA (* p < 0.05, ** p < 0.01, vs WT). (B) Changes in neutralization AUCs (areas under the curves) of SP-swapped vs WT CMU06 with different mAbs. AUC changes are calculated as follows: (SP-swapped AUC – WT AUC). Statistical analysis was performed on neutralization curves (Fig 2A) by 2-way ANOVA (* p < 0.05) (C) ELISA binding of mAbs to gp120 derived from CMU06-WT vs SP-swapped viruses. Virus-derived gp120 was captured by sheep anti-C5 antibody (1μg/ml) and reacted with different mAbs titrated ten-fold from 10 μg/ml. CD4-IgG was titrated five-fold from 10 μg/ml. AUC values were calculated from binding curves. AUC changes relative to the respective WT are shown. Statistical analysis was performed on binding curves by 2-way ANOVA (* p < 0.05) (D) Binding of mAbs to WT vs SP-swapped Envs expressed on the surface of 293T cells. MAbs were tested at following concentrations: 697, 830 and 3685 at 100µg/ml; 2G12 at 50µg/ml; PGDM1400, 2219, 2557, 240D, 98-6 and PGT121 at 25µg/ml; b12 at 20µg/ml; 4E10, 17b at 10µg/ml; PGT128, PGT151, 3BNC117 and CD4-IgG at 5µg/ml; CH59 and NIH45-46 at 2.5µg/ml. Geometric mean fluorescence intensity (MFI) and SD from duplicates in one experiment are shown. Data were analyzed by one-way ANOVA (* p < 0.05, ** p < 0.01, vs WT).

To evaluate the impact of SP swapping on antibody binding, we tested mAb binding to virus-derived gp120 captured onto ELISA plates and native Env expressed on the surface of 293T cells. ELISA with equivalent amounts of Env showed reduced binding of all V2i mAbs tested to gp120 from CMU06-MW, CMU06-398F1, and CMU06-271.1, whereas binding to CMU06-CH119 gp120 was less affected (Fig 2C). Considering that the viruses had an identical V1V2 sequence, the data indicated that SP swapping induced alterations of the V2i structure and/or exposure to reduce mAb recognition. Sporadic changes were also seen with V3 mAbs and CD4-IgG, while no changes were seen with CD4bs mAb NIH45-46.

Next, we tested if SP swapping triggered antigenic changes on Envs expressed on the 293T cell surface. CD4-IgG had comparable binding, indicating equivalent expression of all five Envs. Notably, differences were observed with V2i and V2p mAbs (Fig. 2D). V2i mAb 830 had reduced binding to CMU06-MW, CMU06-398F1, and CMU06-271.1 vs CMU06-WT. Binding of V2i mAb 697 for CMU06-CH119 and CMU06-271.1 was also reduced. V2p mAb CH59 had greater binding to CMU06 with SPs from 398F1, CH119, and 271.1 vs CMU06-WT, and so had V3 crown mAbs 2219 and 2557. gp41-specific mAb (98-6) had reduced reactivity with all SP-swapped Envs vs WT. In contrast, binding of V2q mAb PGDM1400, CD4i mAb 17b, glycan-dependent mAb PGT121 and 2G12, gp120-gp41 interface mAb PGT151, CD4bs mAb NIH45-46, and gp41 mAbs 240D and 4E10 to SP-swapped and WT Envs was comparable. Collectively, these data demonstrated that SP swapping altered Env antigenicity with the most pronounced impact on the V1V2 epitopes.

### Effect of SP swapping on REJO Env incorporation, glycosylation and antibody recognition

To evaluate the effect of SP swapping on a different Env backbone, we tested a clade B tier 2 T/F IMC REJO and swapped its SP with SPs of MW (clade C, chronic, tier 1A), AA05 (clade B, acute), AC02 (clade B, acute), and NL4.3 (clade B, chronic, tier 1) (Fig 3A). SP swapping resulted in viruses with comparable infectivity to REJO-WT, except for REJO-MW which had reduced infectivity (Fig 3B). SPs of AA05, AC02 and NL4.3 increased total Env incorporation, while MW SP lowered Env incorporation. The amount of gp120 (lower band) in REJO-MW was also reduced by 90% (Fig 3C). The poor incorporation of cleaved functional Env into REJO-MW correlated with low infectivity of this virus.

**Figure 3.**
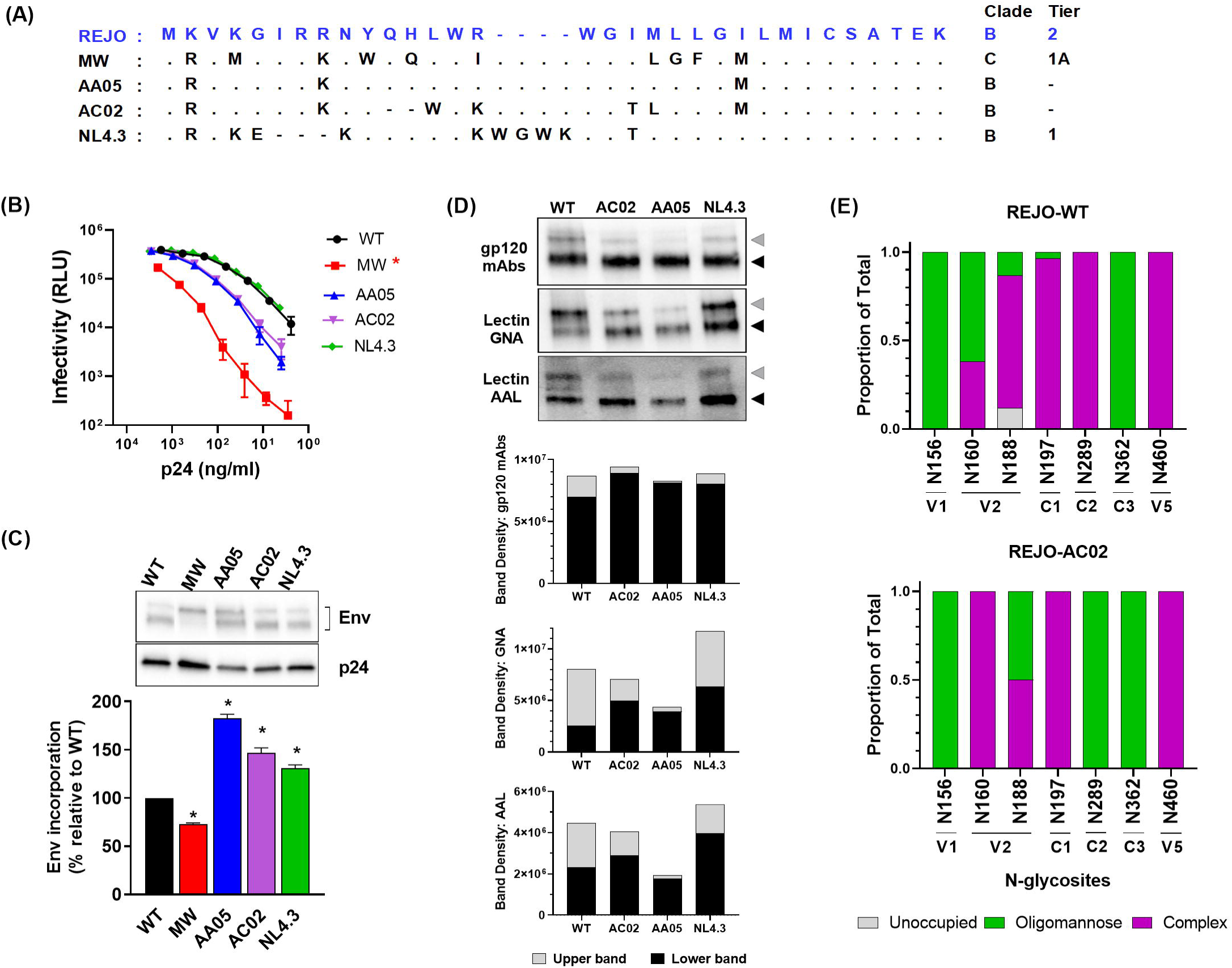
Effects of SP swap on REJO. (A) REJO WT and swapped SPs. Red triangle: SP cleavage site. Within the alignment, dots indicate identity with the WT SP sequence, dash (-) indicates missing residue. (B) Infectivity of WT vs SP-swapped REJO derived from 293T cells in TZM.bl cells. *, p = 0.01 vs WT by Mann-Whitney test. (C) Measurement of Env incorporation into virions by Western blot. *, p≤ 0.005 vs WT by unpaired t-test. (D) Analysis of REJO Env glycosylation by lectin-probed Western blotting. (E) Relative amounts of unoccupied, oligomannose, and complex glycans at seven glycosites detected on both the WT and AC02 SP-swapped REJO Envs as determined by LC-MS/MS.

We subsequently assessed the impact of SP switch on REJO Env glycosylation by lectin-probed Western blotting (Fig 3D). Comparable WT and SP-swapped total Env bands were detected by anti-gp120 mAbs. However, differences were observed with lectins (GNA and AAL) (Fig 3D). Compared with WT, the lower band of REJO-NL4.3 reacted more strongly with GNA (2.5-fold) and AAL (1.7-fold). Both GNA and AAL showed reduced binding to the upper band of REJO-AA05 and increased binding to the lower band of REJO-MW. The LC-MS/MS analysis was able to detect 36% (10/28) and 46% (13/28) of glycosites present on the virion-derived Envs of REJO-WT and REJO-AC02 respectively. Glycan compositions on seven glycosites detected in both Envs are compared (Fig 3E). The data showed altered glycan occupancy at N160 and N188 on the V1V2 apex, as well as at N289 in the C2 region. While 62% of N160 was occupied by oligomannose in REJO-WT, it was 100% decorated with complex-type glycans on REJO-AC02. The oligomannose occupancy of N188 was 13% in REJO-WT but increased to 50% in REJO-AC02. In REJO-WT N289 was 100% oligomannose but became 100% complex in REJO-AC02. The N362 and N460 glycans were not altered. Thus, as seen with CMU06, SP swapping of REJO also altered Env glycan composition with a significant impact on the two V1V2 glycosites detected in the study.

Similar to CMU06, virus sensitivity to neutralization was also altered for SP-swapped REJO vs WT; however, the effect was most pronounced for REJO-AC02, REJO-AA05 and REJO-NL4.3 which became more resistant to most of the V2i mAbs tested (Fig. 4A, B). REJO with AA05, AC02, and NL4.3 SPs also became more resistant to V2q mAb PG9 and to lesser extent PGT145 (Fig. 4A, B). In contrast, the REJO-MW sensitivity to V2i and V2q mAbs was minimally altered. REJO-MW, however, was more sensitive to V3 mAb 2219 and CD4bs mAbs VRC01 and 3BNC117.

**Figure 4.**
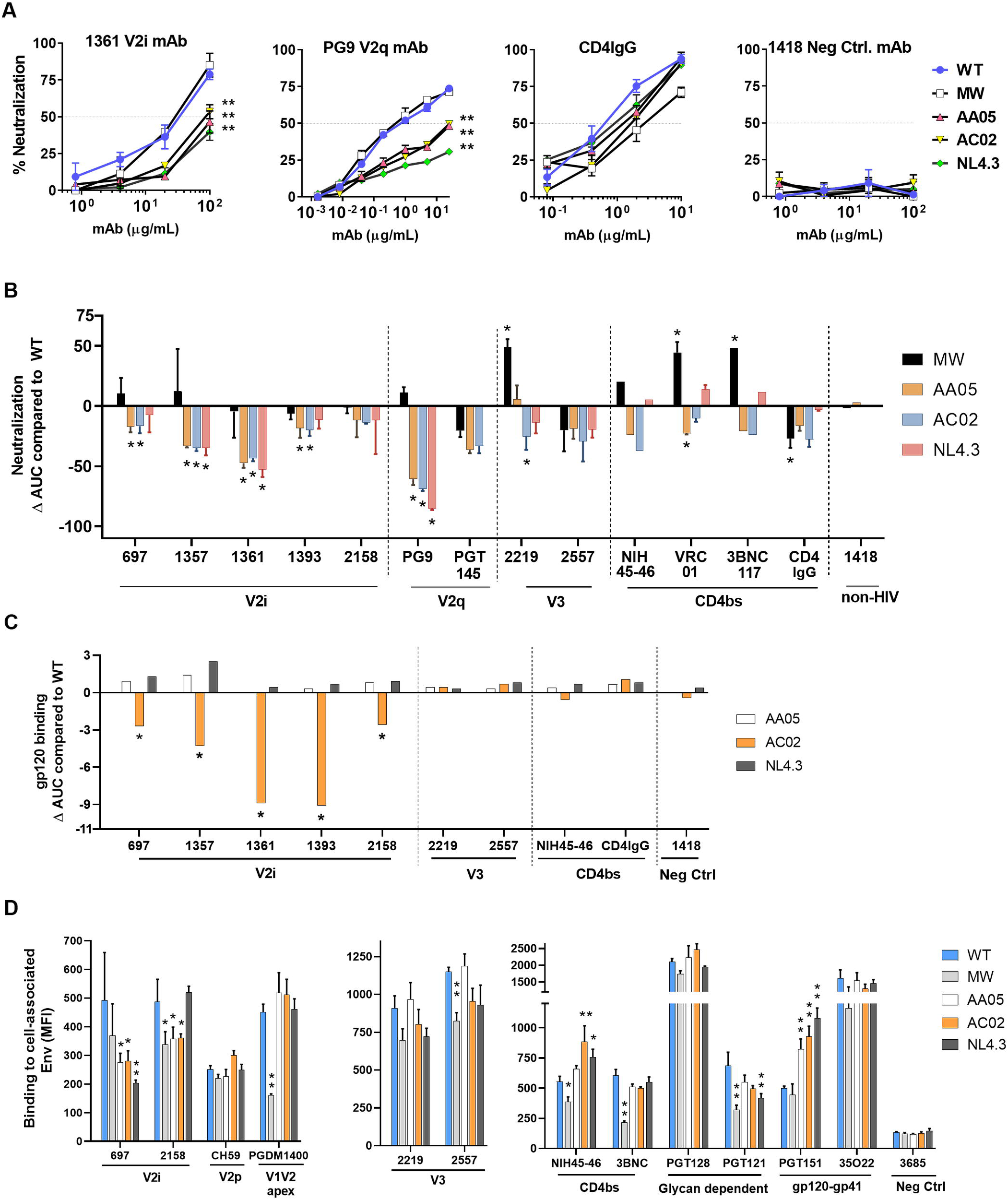
Effects of SP swap on REJO Env-antibody interaction. (A) Neutralization of WT and SP-swapped REJO treated with titrated amounts of anti Env mAbs. Neutralization was performed as in Fig 2A-B. Representative titration curves for V2i mAb 1361, V2q mAb PG9, CD4IgG and negative control mAb 1418 are shown. **, p< 0.001 vs WT by 2-way ANOVA. (B) Changes in neutralization AUCs of WT vs SP-swapped viruses by different mAbs. Statistical analysis was performed on neutralization curves (Fig 2A) by 2-way ANOVA (* p < 0.05) (C) ELISA binding of mAbs and CD4-IgG to gp120 from WT and SP-swapped viruses. Titration was done as in Fig 2C and AUC changes relative to the WT are shown. *, p< 0.05 vs WT by ANOVA of the titration curves. (D) Binding of mAbs to REJO WT vs SP-swapped Envs expressed on the surface of 293T cells. MAbs were tested at the following concentrations: 697, 2158 and 3685 at 100µg/ml; 2219 and 2557 at 50µg/ml; PGT128, PGT121 and 35O22 at 25 µg/ml; CH59, 3BNC117 at 20µg/ml; PGDM1400 and PGT151 at 10µg/ml; NIH45-46 at 2.5µg/ml. Averages and SEM of MFI from duplicates of two experiments are shown. *, p< 0.05; ** p< 0.01 vs WT by one-way ANOVA.

To investigate the effect of SP swaps on REJO Env antigenicity, we assessed mAb binding to virus-derived gp120 (Fig. 4C) and Env expressed on 293T cells (Fig. 4D). Reduced binding of all V2i mAbs was seen with REJO-AC02. In contrast, binding of V3 mAbs, CD4bs mAb NIH45-46, and CD4-IgG to REJO-AC02 was unaltered. REJO-MW was excluded because of low gp120 content. When we examined mAb binding to native Env on 293T cells, a different pattern was observed. Lower binding of V2i mAbs and CD4bs mAbs was observed for one or more SP-swapped Envs. Swapping MW SP onto REJO Env almost abrogated the binding of trimer specific V2q mAb PGDM1400 and reduced the binding of V2i mAbs 697 and 2158, without affecting the binding of V2p mAb CH59. As compared with REJO-WT, REJO-MW also displayed reduced reactivity with CD4bs mAbs NIH45-46 and 3BNC117, and V3 glycan-specific PGT121. In contrast, REJO-AC02, REJO-AA05, and REJO-NL4.3 had lower reactivity mainly with V2i mAbs (697 and/or 2158). PGT151 mAb, which binds specifically to the cleaved Env trimer, showed increased binding to REJO-AC02, REJO-AA05, and REJO-NL4.3, while binding to REJO-WT and REJO-MW was comparable. Hence, like CMU06 SP swapping, REJO SP exchanges also modified mAb interactions with different formats of REJO Env, with the most common effects of reduced mAb recognition against V1V2.

### Strain-specific impact of SP swapping on SF162

To assess the consequences of SP exchanges on another virus strain, we generated SF162 viruses with WT and SP-swapped Envs (Fig. 5A). SF162 is a tier-1A neutralization-sensitive, chronic, clade B isolate. All SP-swapped and WT SF162 viruses displayed comparable levels of infectivity and Env incorporation (Fig. 5B, C). The lectins GNA and AAL reactivity demonstrated moderate changes of oligomannose and fucose-bearing complex glycans on SP-swapped Envs vs WT (Fig. 5D). However, in comparison with CMU06, the same SP exchanges did not make SF162 more resistant to V2i mAbs. The SP-swapped viruses were slightly more sensitive to neutralization by some V2i mAbs and CD4bs mAbs (Fig. 5E). Compared to WT, SF162-MW especially were more sensitive to V2i mAbs 1357 and 1361 and CD4bs mAbs VRC01 and 3BNC117. SF162-398F1 and SF162-CH119, on the other hand, were more resistant to MPER gp41 mAb 2F5. Neutralization by V3 mAbs was not markedly changed.

**Figure 5.**
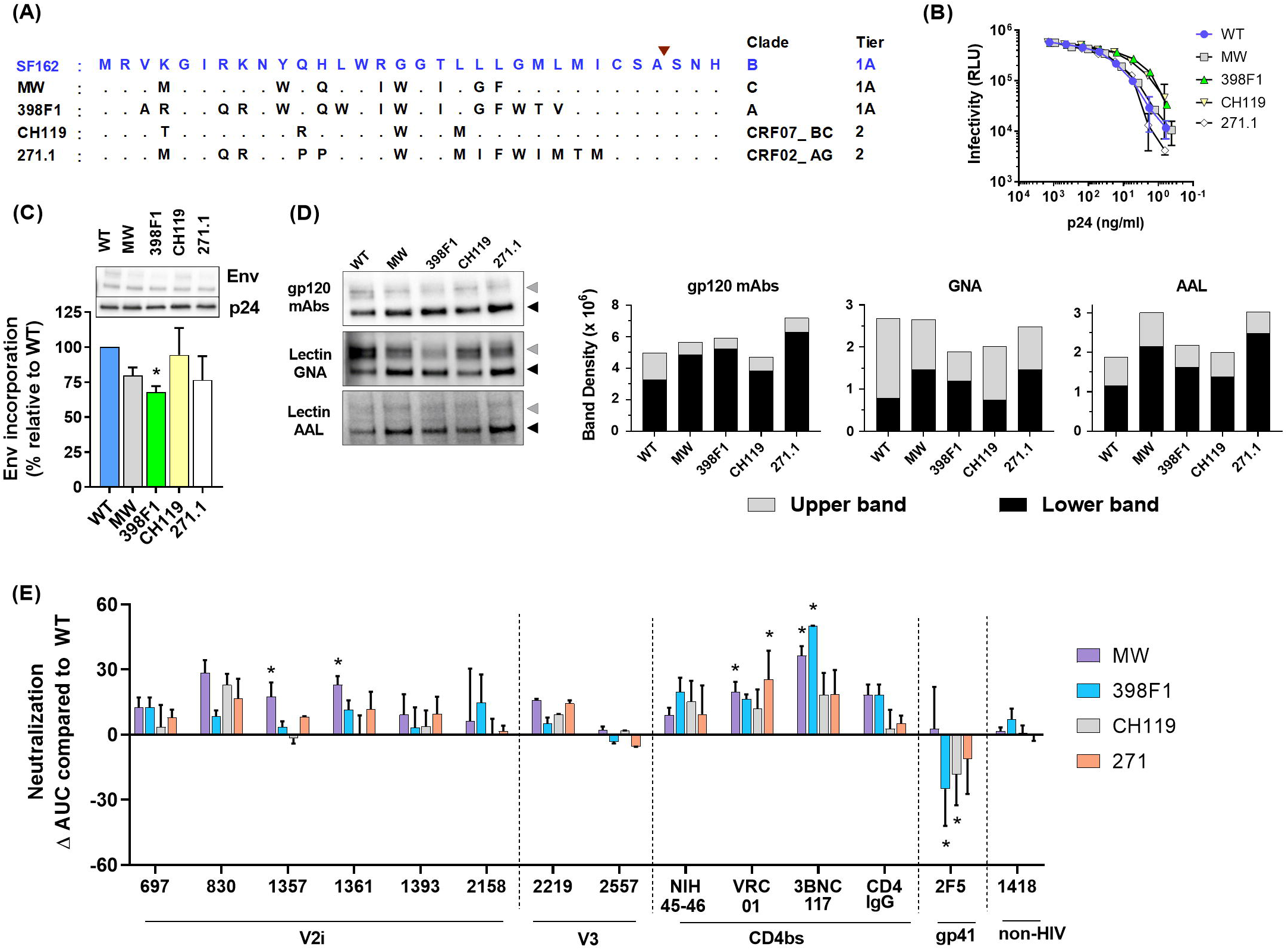
Effects of SP swap on SF162. (A) SF162-WT and swapped SPs. Red triangle: SP cleavage site. Within the alignment, dots indicate identity with the WT SP sequence. (B) Infectivity of 293T-derived SF162 WT vs SP-swapped viruses in TZM.bl cells. (C) Measurement of Env incorporation by Western blot. *, p = 0.017 vs WT by unpaired t-test. (D) Binding of Env from SF162 WT and SP-swapped viruses by gp120 mAbs and lectins (GNA and AAL) in Western blots. Densities of upper and lower Env bands (gray and black arrows) are shown. (E) Changes in neutralization AUCs of WT vs SP-swapped viruses with different mAbs. Statistical analysis was performed on neutralization curves as in Fig 2A by 2-way ANOVA (* p < 0.05)

Figures S4 and S5 summarize the differential effects of SP exchanges on neutralization phenotypes that depend on the Env backbones. These data illuminate the pronounced effects on neutralization by V1V2 mAbs: the SP swaps reduced neutralization sensitivity of CMU06 and REJO but had opposing effects on SF162. MW SP increased neutralization sensitivity to many V2i, V2q, V3, and CD4bs mAbs, which varied depending on the Env backbones.

### Host-cell dependence of SP impact on glycosylation and neutralization

Because glycosylation is host cell-dependent (61, 62), we investigated the effect of SP exchanges on CMU06 produced in primary CD4^+^ T cells. PBMC-derived SP-swapped CMU06 viruses displayed a radically distinct neutralization pattern from 293T-produced counterparts (Fig 6 A-B, Fig S4, S5). While 293T-derived SP-swapped CMU06 acquired increased resistance to V2i mAbs, PBMC-derived viruses with SPs from MW and 398F1 (both tier 1 isolates), but not CH119 and 271.1 (both tier 2), became more sensitive to neutralization by some or all V2i mAbs and to V2p mAb. Remarkably, all PBMC-derived SP-swapped viruses became more resistant to V2q mAbs PG9 and PG16, although neutralization by most CD4bs mAbs and CD4-IgG was largely unaltered. Only swapping with CH119 SP reduced sensitivity to CD4bs mAb VRC01 and MPER gp41 mAb 2F5, without affecting neutralization by V2i and V2p mAbs. V3 crown mAbs had neutralization <50% against CMU06-WT, and the SP-swapped viruses were even more resistant than WT (Fig 6B).

**Figure 6.**
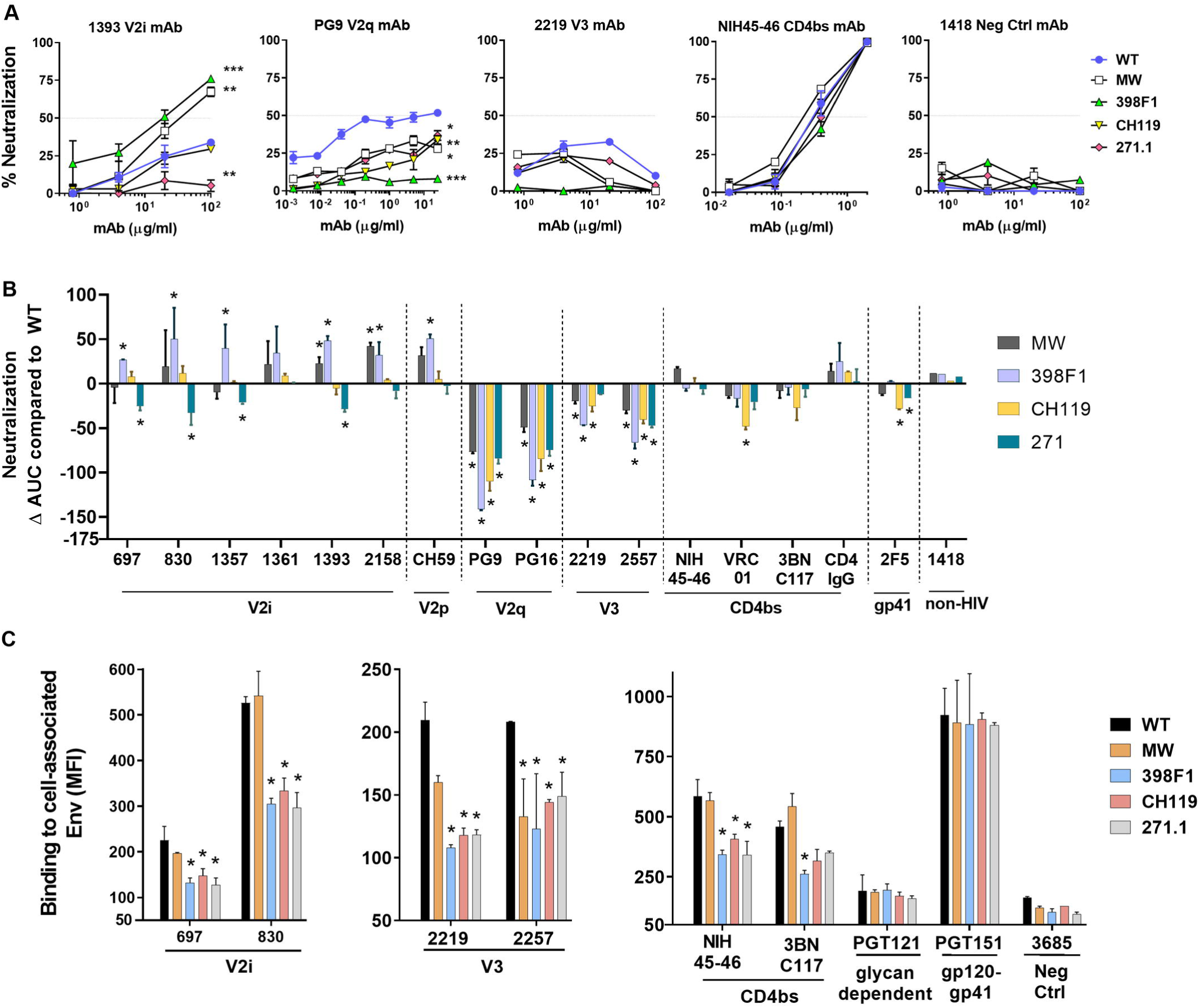
Effect of SP swap on neutralization of CMU06 produced in PBMCs and antigenicity of CMU06 Env expressed on human CD4^+^ T cells. (A) Virus neutralization was measured using TZM.bl cells. MAb 1418 served as a negative control. Neutralization curves of selected virus-mAb pairs are shown. *, p < 0.05, **, p < 0.01, ***, p < 0.001, vs WT by 2-way ANOVA (B) Changes in neutralization AUCs of WT vs SP-swapped viruses by different mAbs. Statistical analysis was performed on neutralization curves as in Fig 2A by 2-way ANOVA (* p < 0.05) (C) Binding of mAbs to Env expressed on CD4^+^ T cell surface. MAbs were tested at the following concentrations: 697, 830, 2219 and 2557 and 3685 at 100µg/ml; PGT121 25µg/ml; CH59 and PGT151 at 10µg/ml; NIH45-46 at 2.5µg/ml; 3BNC117 at 20µg/ml. MFI and SD from duplicates of one experiment are shown. *, p< 0.05 vs WT by ANOVA.

SP swapping also altered the antigenicity of Env expressed on primary CD4^+^ T cells. The changes were more drastic on SP-swapped CMU06 Envs expressed on CD4^+^ T cells (Fig. 6C) vs 293T cells (Fig. 2D). V2i mAbs bound CMU06-398F1, CMU06-CH119, and CMU06-271.1 less than CMU06-WT, but binding to CMU06-MW was unaltered. These SP-swapped viruses also showed reduced binding with V3 mAbs (2219 and 2557), and CD4bs mAbs (NIH45-46 and 3BNC117), although binding of PGT151 and PGT121 was comparable. Collectively, the data show that SP swapping altered the antigenicity of native Env present on virions and cells, albeit in an Env strain- and host cell-dependent manner. Notwithstanding these variabilities, SP swapping clearly affects the V1V2 epitopes.

We further determined glycosylation changes on SP-swapped Envs from PBMC-produced CMU06 using lectin-probed Western blot. Unlike 293T-derived viruses (Fig S2C), we detected predominantly the lower gp120 bands with anti-gp120 mAbs (Fig 7A), suggesting host cell-specific differences in gp120/gp41 processing. Therefore, we analyzed GNA and AAL binding to the lower bands only. Moderate differences were observed in GNA and AAL binding to SP-swapped vs WT CMU06 (Fig 7A). However, the binding patterns were distinct from those seen with 293T-derived CMU06 (S2C Fig).

**Figure 7.**
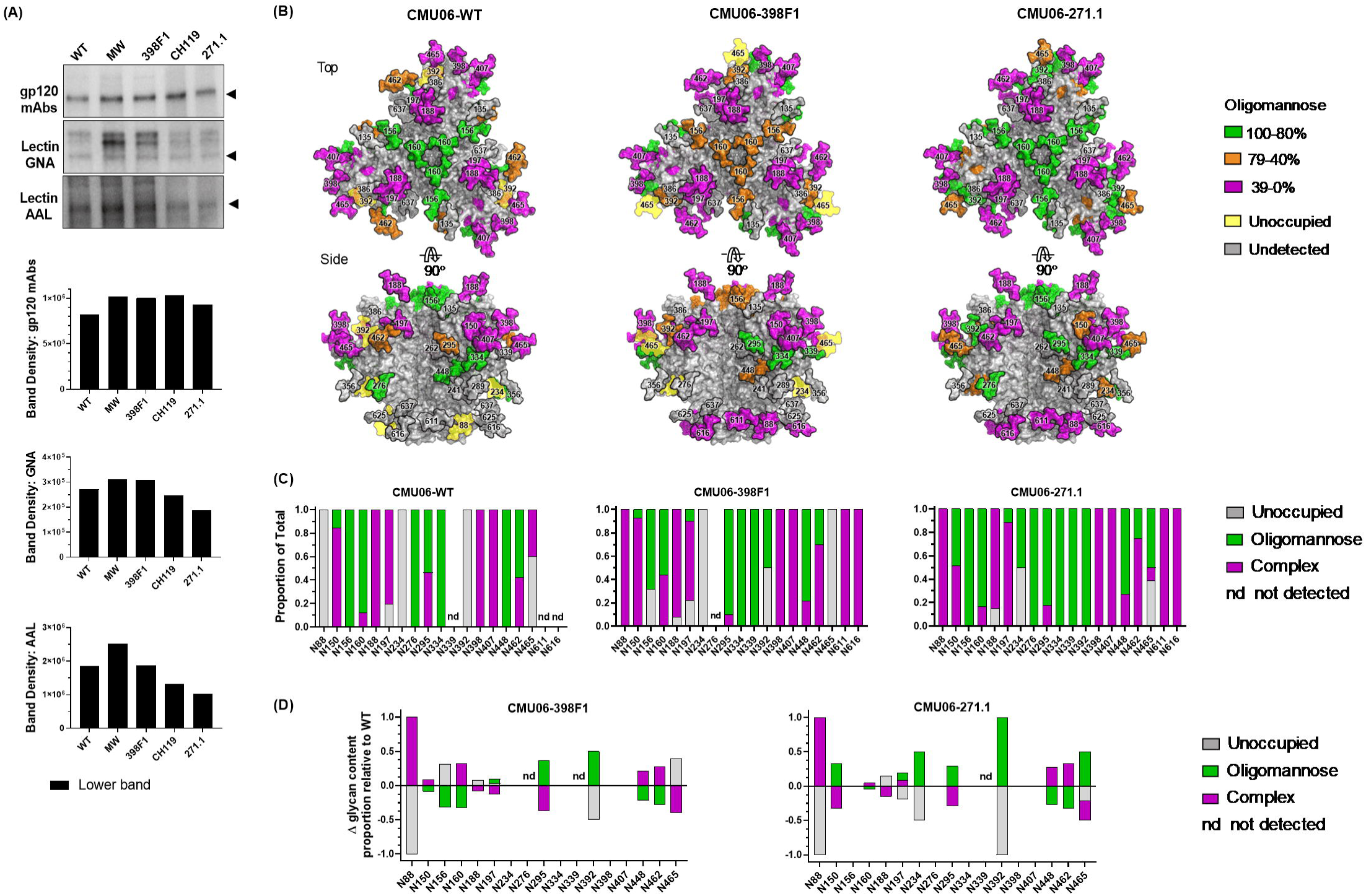
Changes in glycosylation on Env of CMU06 WT and SP-swapped viruses produced in PBMC. (A) Binding of Env by mAbs and lectins (GNA and AAL) measured in Western blots. (B) Models of glycosylated Env trimers from PBMC-derived CMU06 WT and SP-swapped viruses. Models and glycans were generated as in Fig 1B. (C) Proportions of unoccupied, oligomannose, and complex glycans at each glycosite. Glycosites detected in all 3 Envs are shown. nd: not detected in 1 or 2 Envs. (D) Changes in glycan content at each glycosite for CMU06-398F1 and CMU06-271.1 vs CMU06-WT calculated from data in Fig 7C.

We also conducted glycosite-specific analyses of Envs from PBMC-grown viruses by LC-MS/MS (Fig. 7B-D). Of 28 N-glycosites in CMU06 gp160, we detected 16, 18 and 19 in CMU06-WT, CMU06-398F1 and CMU06-271.1, respectively. When the 15 glycosites detected in all three Envs were compared, the percentages of unoccupied glycans was higher in CMU06-WT (25%) and CMU06-398F1 (21%) compared to CMU06-271.1 (7%) (Fig. 7B, S6 Fig). The oligomannose glycans were higher in CMU06-271.1 (48%) compared to CMU06-WT (34%) and CMU06-398F1 (33%). Likewise, total abundance of complex glycans was also altered in CMU06-WT (40%), CMU06-398F1 (47%) and CMU06-271.1 (45%).

Interestingly, the prominent change at N160 on the V1V2 apex was consistently noted in SP-swapped CMU06 produced in PBMCs and 293T cells (Fig. 1 and 7, S6 Fig). N160 had a decreased oligomannose content upon 398F1 SP swapping, confirming that glycan alterations were incurred by 398F1 SP swap regardless of the producer cells.

We also assessed glycans that were affected by both host cells and SPs; these comprised most N-glycans detected: N88, N150, N156, N188, N197, N295, N392, N407, N448, N462, N465, N611 and N616. Glycan content at these glycosites differed for PBMC- vs 293T-produced viruses and among the three SP-swapped viruses (S6 Fig). Notably, N156 on the V1V2 apex displayed only oligomannose glycans in PBMC-derived CMU06-WT, but had a mix of oligomannose and complex glycans in 293T-derived CMU06-WT. On PBMC-derived CMU06-271.1, N156 also was completely occupied with oligomannose, whereas on 293T-derived counterpart it was mostly unoccupied. N156 had reduced oligomannose contents on CMU06-398F1 vs CMU06-WT, when produced in PBMC or 293T cells. Another example is illustrated by N88 on the gp120-gp41 interface, which was completely unoccupied in PBMC-derived CMU06-WT, but 82% unoccupied in 293T-derived WT. In contrast, N88 was populated exclusively by complex glycans in CMU06-398F1 and CMU06-271.1 from PBMC, although it was 50% and 80% unoccupied in 293T-derived counterparts.

The N398 glycan changes depended on host cells only and are independent of SPs. N398 was 100% occupied by complex glycans on all three PBMC-derived viruses, these were reduced to 22-36% in 293T-derived viruses.

Few glycosites were not identified in all samples, allowing only partial comparison. For example, N334 had only oligomannose glycans on all PBMC-derived viruses and 293T-derived CMU06-398-F1. N611 on gp41 was detected in two PBMC-derived and three 293T-derived viruses and displayed exclusively or mostly complex glycans. N616 had complex glycans on four of five PBMC- and 293T-derived viruses but a mix of complex and oligomannose glycans on CMU06-271.1 from 293T cells.

Collectively, these data revealed the impact of SP changes on glycan compositions at specific glycosites and demonstrated the distinctive glycan profiles of viruses from PBMCs vs 293T cells. Because we compared viruses with identical Env sequences, the observed differences reflect how glycosylation proceeds at each glycosite in the context of different SPs and host cells.

## Discussion

HIV-1 Env glycosylation is influenced by viral and cellular factors. This study demonstrates the effect of Env SP, a highly variable albeit understudied viral element, on glycosylation and consequently on Env properties and virus phenotypes. Introducing SPs from different virus strains to the same Env backbone altered proportions of oligomannose, complex, and unoccupied glycans on multiple glycosites. Interestingly, exchanging CMU06 SP with 398F1 SP reduced the oligomannose content of N156 and N160 glycosites on V1V2 at the Env spike apex essential for the V2q-class bNAbs. This effect was observed in both PBMC and 293T-produced CMU06-398F1 viruses. Swapping the AC02 SP onto the REJO Env similarly reduced the oligomannose content of N160 from 62% on WT to 0% on REJO-AC02. Consistent with glycan changes, these SP exchanges affected Env recognition and virus neutralization by mAbs targeting V1V2 epitopes. Thus, although SP is not a part of the mature Env, it is a determinant that affects Env glycosylation and antibody recognition.

Among the three classes of V1V2 mAbs, V2i and V2q mAbs were prominently affected by SP swaps, as evidenced by increased or reduced resistance of many SP-swapped tier-2 CMU06 and REJO vs their respective WT to neutralization by these mAbs. The effect nonetheless varied depending on SPs and host cells producing the virus. Reasons for altered sensitivity to V2i mAb-mediated neutralization are not fully understood. The V2i epitopes are often occluded on native pre-fusion trimers of tier-2 viruses (22) and become exposed to mAbs only after prolonged virus-antibody incubation (14). The structural dynamics influencing V2i epitope exposure is governed partly by Env glycan composition. Our previous studies showed that enrichment of high-mannose glycans, as detected by lectin western blotting, increased virus resistance to V2i mAbs, even when the mAb-virus incubation was extended to >18 hours (34). Here, using LC-MS/MS approach we observed that the total oligomannose content was increased by 4.4% and 7.9% on 293T-derived CMU06-398F1 and CMU06-271.1, respectively, over that of CMU06-WT. The oligomannose content also increased by 15.5% on PBMC-derived CMU06-271.1. Correspondingly, these SP-swapped viruses showed a trend of or a significant increase in V2i mAb resistance vs WT: the cumulative neutralization AUC values decreased by 87 and 188 for 293T-derived CMU06-398F1 and CMU06-271.1, respectively, and by 125 for PBMC-derived CMU06-271.1. In contrast, the oligomannose content of PBMC-derived CMU06-398F1 was reduced by 6%, and the virus was more sensitive to V2i mAbs (cumulative AUC of 483 vs 253 for WT). These data suggest that increasing oligomannose glycans is one potential strategy that HIV-1 can employ to resist neutralizing activities of V2i Abs, and such alteration can be triggered by changes in Env SP. Consistent with this, a single substitution in the N-terminal SP region was found sufficient to affect Env glycan composition and increase virus resistance to neutralization by V2i mAbs (34).

Unlike V2i mAbs, V2q mAbs PG9 and PG16 directly target oligosaccharide moieties as part of their epitopes (63). Hybrid glycans bearing high mannose and biantennary motifs with two terminal sialic acids at N156 and N160 on the V1V2 apex are important for PG9 binding, whereas PG16 recognizes α-2,6-sialylated hybrid and complex glycans on these sites (20, 64). Sensitivity of 293T-derived CMU06 to PG9 and PG16 was enhanced when CMU06-WT SP was exchanged to MW, 398F1, or 271.1 SP. In contrast, SP-swapped viruses were more resistant to V2q mAbs when produced in PBMCs. Our LC-MS/MS data offered evidence for the first time that the modulation of virus sensitivity to these V2q mAbs correlated with changes in the complex glycan content on the viruses. A decrease of complex glycans overall and at N156 and N160 collectively was observed in 293T-derived CMU06-398F1 and CMU06-271.1, both of which were more sensitive to V2q mAbs as compared to CMU06-WT. Conversely, the total complex glycan content and the proportion of complex glycans at N160 and N156 in combination increased when these viruses were grown in PBMC and the viruses became more resistant to V2q mAbs. Alteration of glycan content was similarly observed for REJO-WT and REJO-AC02, in which total complex glycans on N156 and N160 were higher on REJO-AC02 vs REJO-WT and correlated with increased resistance of REJO-AC02 to neutralization by the V2q mAb PG9. These data indicate that an increased amount of complex glycans is unfavorable for V2q mAbs. Alternatively, the shift may supplant the hybrid-type glycan structures with terminal moieties preferred by these V2q mAbs (20, 64), but cannot be resolved by LC-MS/MS. Glycans proximal to N156 and N160, such as N136 and N149 (undetected here) also have been shown to influence the binding and neutralization by V2q mAbs (20, 65). While more detailed studies are required to define Env glycomes compatible with V2q mAbs, the data clearly shows that the glycan compositions of N156 and N160 are influenced by the SP sequences.

In contrast to V2i and V2q mAbs, mAbs against other Env regions were less affected by SP swapping. Env binding and virus neutralization by CD4-IgG were likewise unperturbed, indicative of the proper folding of all tested SP-swapped Envs. The SP-swapped Envs also maintained the features characteristic of the native trimeric Env, such as the presence of TAMP, which is absent on uncleaved or incorrectly assembled trimers (66-68), and the exclusive oligomannose content at N276 when this glycosite was detected (PBMC-produced CMU06-WT and CU06-271.1). This contrasted to the N276 occupation by complex glycans on BG505 pseudotrimers and gp120 monomers (66). Further, we observed no drastic reduction of PGT151 mAb binding to SP-swapped vs WT Envs, and three SP swaps even increased PGT151 binding to REJO Env. PGT151, which targets glycan-bearing epitopes in the gp120-gp41 interface like mAb 35O22, binds only to properly folded, cleaved trimers (69, 70). Similarly, Env binding and virus neutralization by trimer-dependent V2q mAbs PGT145 and PGDM1400 were largely undisturbed by SP exchanges, except for REJO-MW. We also observed only minimal sporadic effects on neutralization by V3 crown mAbs, demonstrating that SP swapping did not shift Env trimers toward more open conformations (71-73). Altogether, these data indicate that SP swapping subtly modifies the proportion of Env oligosaccharide content and the antigenicity of V1V2 without triggering global structural transformations on Env.

As indicated by comparable Env incorporation and virus infectivity, SP exchanges had no deleterious effects on Env functions. One exception was REJO-MW, which diminished expression of functional cleaved Env gp120, reduced Env incorporation into virions, and decreased virus infectivity. However, this effect was not seen when MW SP was introduced to CMU06 and SF162. The reasons for REJO-MW incompatibility remains unclear. A reduction of cleaved gp120 was previously noted when a mutation was introduced to position 12 of REJO SP (H12Q), but not to that of JRFL SP (Y12Q) (34). Glutamine occupies position 12 of MW SP and not the other SPs tested with REJO, but MW SP also differs from REJO SP at 9 other positions, requiring systematic evaluations for the contribution of glutamine at position 12 vs other MW SP residues.

Glycosylation begins in the ER with the addition of Glc3Man9GlcNAc2 onto asparagine in the NX(T/S) (X≠Proline) sequon on a nascent protein. As the protein crosses the ER and the Golgi apparatus, the high-mannose structure is trimmed and subsequently elaborated with hybrid- and complex-type glycans. In view of the observed effect of SP swapping, we suggest that the extent to which the glycan on a particular glycosite is processed is decided while Env is an ER resident with its SP still tethered. SP may subtly affect the compactness of Env folding, which consequently imposes or releases structural constrains to enzymes that generate hybrid or complex glycans in the Golgi. Nonetheless, the steps at which the SP sequence dictates the glycan content of Env are unknown. The Env SP is cleaved post-translationally and this delayed SP cleavage is seen across HIV-1 subtypes ensuring low Env expression on virions (5, 6, 74). In addition to the positively charged n-region, residues in the hydrophobic SP region (h-region) and downstream of the SP cleavage site are important for this property (5, 6). The h-region forms an alpha-helix across the ER membrane and covers the cleavage site to delay SP cleavage (3, 5). Since the cleavage sites in SP-swapped Envs and WT counterparts are identical, perhaps the h-region variations affect the cleavage site accessibility for signal peptidase, altering the efficiency of SP cleavage of Env with heterologous SPs, similar to that seen with the h-region-mediated regulation of SP cleavage for preprolactin (75, 76).

Prediction based on Model SignalP 3.0 (77) indicates that the cleavage probabilities were slightly increased for CMU06 with 398F1, CH119, and 271.1 SPs vs WT (S7 A-B Fig), with 271.1 SP having the highest probability. Topology and structure predictions using PSIRED (78, 79) further show differences in the cytoplasmic and ER regions and in the alpha-helical structure around the cleavage site (S7C-D Fig). The transmembrane-spanning residues encompass positions 16-31 for CMU06 WT and CMU06-CH119 but differ for the other three swaps. The 271.1 SP cleavage site is also shown to be most exposed (S7D Fig), but experimental data are needed to verify if cleavage efficiency is indeed changed to affect the rate of Env synthesis. Past studies demonstrated, however, that replacing SPs of influenza hemagglutinin and HIV-1 Env by tPA/22P SP, which increased SP cleavage probability to 0.925, caused higher secretion of these proteins (80). Similarly, M26P mutation in HIV-1 HXB2 SP was found to increase the SP cleavage probability from 0.628 to 0.928 and change the SP cleavage from post-translational to co-translational (6). Prolines are notably absent near SP cleavage sites in 99.9% of HIV-1 gp160 (6).

In additional to SP cleavage, SP interaction with the signal-recognition particle (SRP) may also play a role in determining Env phenotypes. The h- and n-regions, which vary greatly among SPs in length and sequence, are deemed important for SRP interaction (81-83). Thus, it is possible that different SPs may bind SRP with different affinities impacting the chaperone-mediated Sec61 channel gating and the interaction with chaperones involved in protein folding and assembly (75, 84, 85). A single substitution of a helix-breaking glycine residue by a helix-promoting leucine residue in the hydrophobic core of prokaryotic SPs can promote SRP binding (86, 87) and regulate the pathway of protein translocation (88). The SPs studied here differ in the number of basic, hydrophobic, and leucine residues (S1 Fig); the significance of each element needs to be better understood, considering that SP exchange is a common strategy to promote secretion of recombinant Env vaccines and SP selection is critical to generate glycomes faithfully representing those of native Envs.

In summary, this study demonstrates an important role of HIV-1 Env SP in influencing Env glycan content. By introducing SP from a particular HIV-1 strain, we can modulate the relative proportion of unoccupied, high-mannose, and complex glycans on specific glycosites, including N156 and N160 on V1V2 at the apex of Env spike. SP swaps also can alter Env recognition and virus neutralization by mAbs, especially mAbs against V1V2. Data from this study have significant implications for vaccine development: SP is a critical component that must be rationally selected and incorporated into the design of Env-based HIV-1 vaccine.

## Materials and methods

### Plasmids

The infectious molecular clones (IMC) were generated by cloning the CMU06 and SF162 Envs into pNL4.3 backbone to construct pNL-CMU06 and pNL-SF162, respectively. To generate SP-swapped CMU06 IMCs, restriction site NheI was introduced at positions 89-90 (CGGAGT to GCTAGC) without amino acid alterations (A29, S30). DNA fragments with swapped SPs were synthesized (Genscript) and digested using EcoRI (in pNL4.3 backbone) and NheI and ligated to pNL-SF162 and pNL-CMU06 digested with the same enzymes. The pREJO.c/2864 IMC (REJO) was used to make REJO SP-swapped plasmids by a multi-step overlapping PCR mutagenesis strategy using Pfx50™ DNA Polymerase PCR System (34). Briefly, in the first PCR step, mutated fragments were individually generated in two separate reactions. The primer pairs AvrIIF/MWR and MWF/BstEIIR were used to generate pREJO-MW (STAR Methods). The PCR fragments were agarose gel-purified, combined, and added to a second-stage PCR with the flanking primers AvrIIF and BstEIIR. Products of the second-stage PCR were digested by AvrII and BstEII restriction enzymes and inserted into the AvrII- and BstEII-digested fragment of pREJO.c/2864 to yield REJO-MW. The other REJO plasmids were constructed similarly using the primers listed in STAR Methods. All the plasmids were sequenced to confirm the presence of the desired sequence changes without any other mutations.

### Cell lines

HEK293T/17 cells (293T) were used to produce infectious HIV-1 viruses. TZM.bl cell line was used to assay virus infectivity and virus neutralization. TZM.bl cell line is derived from HeLa cells and genetically modified to express high levels of CD4, CCR5 and CXCR4 and contain reporter cassettes of luciferase and β-galactosidase that are each expressed from an HIV-1 LTR. The 293T and TZM.bl cell lines were routinely subcultured every 3 to 4 days by trypsinization and were maintained in Dulbecco’s modified Eagle’s medium (DMEM) supplemented with 10% heat-inactivated fetal bovine serum (FBS), HEPES pH 7.4 (10 mM), L-glutamine (2 mM), penicillin (100 U/ml), and streptomycin (100 μg/ml) at 37°C in a humidified atmosphere with 5% CO2.

### Viruses

Infectious viruses were generated by transfecting 293T cells with wild type (WT) or SP-swapped pNL-CMU06, pREJO and pNL-SF162 plasmids using jetPEI transfection reagent (34). Supernatants were harvested after 48 hours and clarified by centrifugation and 0.45μm filtration. Single-use aliquots were stored at −80°C until use. Virus infectivity was assessed on TZM.bl cells as described (34, 89). Briefly, serial two-fold dilutions of virus stock in 10% DMEM were incubated with TZM.bl cells (in duplicates for each dilution) in half-area 96-well plate in presence of DEAE-Dextran (12.5 µg/ml) for 48 hours at 37°C. Virus infectivity was measured by β-galactosidase activity. The dilution of a virus stock that yielded RLUs between 150,000 and 200,000 was used in neutralization assays, as detailed below. HIV-1 p24 ELISA assay was used to quantify the p24 content in each virus stock using the manufacturer’s protocols.

To generate PBMC-derived viral stocks, PBMC were isolated by lymphoprep gradient centrifugation from leukopaks of HIV-1 seronegative donors. Prior to HIV-1 infection, PBMC were activated by incubation in RPMI-intereukin-2 (IL-2) growth medium containing 10 μg of phytohemagglutinin (PHA) (PHA-P) per ml. The RPMI-IL-2 growth medium was RPMI 1640 medium supplemented with 10% heat-inactivated fetal calf serum, L-glutamine (2 mM), penicillin (100 U/ml) and streptomycin (100 μg/ml) and 20 U of recombinant IL-2 per ml. After overnight incubation with PHA, cells were washed and cultured with IL-2 for additional 2 to 3 days. All cultures were maintained in 5% CO2 incubators at 37°C. PBMCs were exposed to virus (50ng p24 of 293T-derived stock) overnight and then washed to remove the viral inoculum. Virus-containing supernatant was harvested on day 7 and day 10. Virus infectivity and p24 contents were measured as above.

### Human monoclonal antibodies

V2i, V3, and control mAbs were produced in our laboratory as described (26, 90-96). Other mAbs used in this study were obtained through the NIH AIDS Reagent Program, Division of AIDS, NIAID, NIH. MAbs specific to different Env epitopes were tested, including: V2i (697, 1357, 1361, 1393 and 2158), V2p (CH59), V2q (PG9, PG16, PGT145 and PGDM1400), V3 (2219, 2557, 3074 and 3869), CD4bs (NIH45-46, VRC01, b12, 3BNC117), CD4i (17b), glycan dependent (PGT121, PGT128 and 2G12), gp41 (240-D, 98-8, 2F5 and 4E10), and gp120-gp41 interface (PGT151 and 35O22). An CD4-IgG fusion protein was also included, along with an irrelevant anti-parvovirus mAb 1418 and anti-anthrax mAbs 3685 used as negative controls.

A cocktail of anti-human anti-gp120 mAbs (anti-V3: 391, 694, 2219, 2558; anti-C2: 846, 1006; anti-C5: 450, 670, 722, 1µg/ml each) were used to detect Env. The mAb 91-5D (1µg/ml) was used to detect Gag p24.

### Virus neutralization assay

Neutralization activity of HIV-1-specific antibodies was measured as a reduction in β-galactosidase reporter gene expression of TZM-bl cells after infection of TZM-bl cells with WT and SP-swapped viruses. Neutralization was performed using either the standard 1 hour assay (14, 34, 97) or the prolonged 24 hour assay (14, 97). Serially diluted antibodies and viruses (150,000-200,000 RLUs) were pre-incubated for 1 hour or 24 hours prior to adding the target TZM.bl cells. After a 48-hour incubation at 37°C and 5% CO2 incubator, the β-galactosidase activity was measured. Each condition was tested in duplicate or triplicate, and experiments were repeated 2-4 times. Percent neutralization was determined based on virus control (TZM.bl cells with virus alone) and cell control (TZM.bl cells only) under the specific assay condition. Neutralization assays were performed after a 24-hour mAbs-virus incubation for all mAbs except PG9, PGT145, NIH45-46, VRC01, 3BNC117, and CD4-IgG which were pre-incubated with viruses for 1 hour.

### Western blot analyses with antibodies and lectin probes

Western blot analyses were performed to quantify the ratios of Env to p24 proteins incorporated into the WT and SP-swapped viruses and to evaluate Env reactivity with different lectins as in (34). The virus particles sucrose-pelleted from supernatants were lysed in SDS-PAGE loading buffer, resolved on 4–20% tris-glycine gels, and blotted onto nitrocellulose membranes, blocked overnight with 5% skim milk powder in phosphate buffered saline (PBS) followed by probing with antibodies or lectins. A cocktail of anti-human anti-gp120 mAbs (anti-V3: 391, 694, 2219, 2558; anti-C2: 846, 1006; anti-C5: 450, 670, 722, 1µg/ml each) were used to detect Env. The mAb 91-5D (1µg/ml) was used to detect Gag p24. Biotinylated GNA, and biotinylated AAL were each used at 2µg/ml. Lectin binding was detected with HRP-neutravidin (1:1500 for 1 hr RT). All dilutions were made in Superblock T20 Buffer. Membranes were developed with Clarity Western ECL substrate and scanned by ChemiDoc Imaging Systems (Bio-Rad Laboratories). Purified recombinant gp120 and p24 proteins were also loaded at a known concentration as controls (data not shown). Band intensities were quantified using the Image Lab Software Version 5.0.

### gp120 – mAb binding assay

The relative binding of mAbs to gp120 from WT and SP-swapped viruses was measured by a sandwich ELISA. Half-area high-binding ELISA plates were coated with sheep anti-C terminal gp120 Abs (1µg/ml in PBS), blocked with 2% bovine serum albumin (BSA) in PBS, and incubated with 1% Triton X100-virus lysates containing 20 ng/ml Env (quantitated by Western blots). Serially diluted mAbs (0.01-10μg/ml) were then added for 2 hours, and the bound mAbs were detected with alkaline phosphatase-conjugated goat anti-human IgG and p-nitrophenyl phosphate substrate. The optical density (OD) was read at 405nm using BioTek PowerWave HT Microplate Spectrophotometer.

### Cell-associated Env binding assay

Assay to detect antibody binding to cell surface-expressed Env was performed as described (98) with minor modifications. A total of 4 × 10^6^ 293T cells were seeded in 15 ml of culture medium in a 100-mm tissue culture dish and incubated at 37 °C. Twenty four hours later, cells in each dish were transfected with 20 μg gp160 expression plasmid (WT or SP swaps) using JetPEI (DNA:JetPEI ratio of 1:3) following manufacturer’s instructions. The transfected cells were incubated for 24 hours at 37 °C, washed with PBS, detached with trypsin-free cell-dissociation buffer, and resuspended in PBS containing 2% BSA. Cells were stained with Live/dead Aqua stain and distributed into 96-well round-bottom tissue culture plates (5×10^4^/well) for individual staining reactions. Cells were incubated with mAbs at concentrations detailed in figure legends. For detection of mAb binding, biotinylated goat anti-Human IgG Fc (1:1000) followed by streptavidin phycoerythrin (PE) (1:500) was used. The cells were washed 3X with PBS-B (PBS plus 1%BSA) after each step and all incubation steps were performed on ice for 30 min. Cells were analyzed with a BD Fortessa flow cytometer, and 30,000 events were collected in the PE+ gate. Analysis was carried out using FCS-Express software as follows: 293T cells were selected from a plot of forward-area vs. side scatter-area (FSC-A/SSC-A) from which doublets were excluded in a forward scatter height vs forward scatter area plot (FSC-H/FSC-A). Live cells were selected by Aqua-negative gating, and geometric mean fluorescent intensity (MFI) of PE+ cells, representing anti-Env-stained cells, were quantified. Background MFI, as determined from cells stained without primary antibodies was subtracted from all Env-mAb pairs.

For testing mAb binding to Env expressed on primary CD4+ T cells, the experiments were conducted as above with the following modifications: CD4+ T cells were isolated from PBMCs (isolated from Leukopaks) using an EasySep Human CD4+ T cell Enrichment Kit. PHA-activated CD4+ T cells were infected with WT or SP-swapped viruses (500 ng p24/million cells) by spinoculation at 1200×g for 2 hours. The virus inoculum was replaced with fresh medium (RPMI 1640 media supplemented with 10% FBS with 20 U/mL IL-2) and the cells were further incubated for 7 days at 37°C. Cells were harvested for mAb staining as above.

### Mass Spectrometry

Analysis for site-specific glycosylation was performed as in (99). Briefly, 293T- and PBMC-derived infectious virus stocks were concentrated by sucrose cushion centrifugation. Concentrated (250X) virus preparations were loaded on SDS-PAGE gel (7.5%) and the separated Env bands were excised to use directly for mass spectrometry (MS).

The SDS-PAGE gel bands were washed with 100% acetonitrile and water three times. A proteomics-based strategy was used to assess the degree of glycan processing and the degree of site-occupancy of each glycosite (99). The proteins in the gel bands were denatured and alkylated by 10 mM dithiothreitol (DTT) and 55 mM iodoacetamide in 25 mM ammonium bicarbonate, respectively. The resulting proteins were digested with the combination of trypsin and chymotrypsin at an enzyme/substrate ratios of 1:15 (w/w) and 1:10 (w/w) respectively in 25 mM ammonium bicarbonate. Sequential treatment with two endoglycosidases was then performed to introduce novel mass signatures for peptides that contain glycans of high-mannose types and complex-type glycans (99). First, the Env peptides were digested with Endo H to cleave high-mannose (and hybrid) glycans between the innermost GlcNAc residues, leaving a GlcNAc attached to the Asn (N+203). The subsequent PNGase F treatment removed the remaining complex-type glycans, and in the process converted Asn to Asp, resulting in a +0.984 Da mass shift (N+1) (100, 101). For peptides with unoccupied glycosites, these treatments produce no mass shift (N+0). Using this strategy, liquid chromatography–mass spectrometry (LC-MS/MS) data were acquired for each sample. Peptides were identified using SEQUEST. The abundance of each peptide was determined by the sum of the peak areas from all identified charge states (63).

The 293T-derived Env samples were analyzed on a Q-Exactive mass spectrometer. De-glycosylated peptides were separated on a Dionex Ultimate 3000 RSLC nano system with a 75 µm × 15 cm Acclaim PepMap100 separating column protected by a 2-cm guard column. The flow rate was set at 300 nl/min. Buffer A and B were 3% ACN (0.1% FA) and 90% ACN (0.1% FA), respectively. A 130-minute gradient was run, consisting of the following steps: 0-10 min, 2-5% B; 10-90 min, 5-25% B; 90-112 min, 25-35% B; 112-115 min, 35-95%; 115-125 min, hold at 95% B; 125-129 min, 95-2% B; 129-130 min, hold at 2% B. Orbitrap MS1 spectra (AGC 3e6) were collected from 400– 1,800 m/z at a resolution of 70 K followed by data dependent higher-energy collisional dissociation tandem mass spectrometry (HCD MS/MS) (resolution 35,000 and collision energy 31%) of the 12 most abundant ions using an isolation width of 1.4 Da. Dynamic exclusion time was set at 30s.

The PBMC-derived Env samples were analyzed on an Orbitrap Fusion Lumos tribrid mass spectrometer. Approximately 1 µg of de-glycosylated peptides were injected directly onto a 28 cm, 75 um ID column packed with 1.9 um Reprosil-Pur C18-AQ beads (Dr. Maisch GmbH). The flow rate was separated at 300 nl/min on an EASY-nLC 1200. Buffer A and B were 3% ACN (0.1% FA) and 90% ACN (0.1% FA), respectively. A 110-minute gradient was run, consisting of the following steps: 0-1 min, 2-6% B; 1-85 min, 6-30% B; 85-94 min, 30-60% B; 94-95 min, 60-90% B; 95-100 min, hold at 90% B; 100-101 min, 90-50% B; 101-110 min hold at 50% B. Peptides were eluted directly from the tip of the column and nanosprayed into the mass spectrometer. The Orbitrap Fusion Lumos was operated in a data-dependent mode. Full MS1 scans were collected in the Orbitrap at 60 K resolution with a mass range of 350–2,000 m/z and an AGC target of 4e5. The cycle time was set to 2 s, and within this 2 s the most abundant ions per scan were selected for CID MS/MS in the orbitrap with an AGC target of 5e4. Maximum injection times were set to 50 ms for both MS and MS/MS scans. Monoisotopic precursor selection was enabled, and dynamic exclusion was used with exclusion duration of 45 s.

CMU06 Env trimer model was generated using a SOSIP.664 trimer (PDB 5FYJ) at each glycosite, except N651 due to lack of template. Glycan was added onto models based on the LC-MS/MS data of virion-associated Envs from 293T cells and PBMC. For sites with high complex-type content, biantennary glycan with core fucose Gal2GlcNAc2Man3FucGlcNAc2 moieties were used. For those with high oligomannose content, Man5GlcNAc2 was added. Protein surface and glycans that were undetermined or absent on CMU06 were colored gray. Glycosites at N135, N234, N241, N262, N276, N289, N356, N386 and N625 were undetected in all Envs and are not shown.

CMU06 Env trimer protein model was generated using a SOSIP.664 trimer (PDB 5FYJ). Glycans were added onto the model based on the LC-MS/MS data of virion-associated Envs from 293T cells except glycan N674 due to the lack of a template. For glycosites with high complex-type content, biantennary glycans with core fucose Gal2GlcNAc2Man3FucGlcNAc2 moieties were used. For those with high oligomannose content or undetected, Man5GlcNAc2 were added. Glycans that were undetected (e.g., N135, N234, N241, N262, N276, N289, N356, N386 and N625) are colored gray, while those that were unoccupied on PBMC-derived CMU06 are displayed by semi-transparent yellow patches.

## Statistical analysis

Comparison of SP swaps vs WT groups and controls was done using analysis of variance (ANOVA) or by Student’s t-test as mentioned in the figure legends. Statistical analyses were performed with GraphPad Prism 8.

## Acknowledgements

The authors thank Ms. Vincenza Itri and Ms. Xiaomei Liu for providing HIV-1-specific mAbs and Ms. Alisa Fox for assistance with flow cytometry. This study was supported in part by NIH R01 grant (AI140909 to C.U.), NIH R21 grant (AI124863 to C.U. and C.E.H.), NIH R01 (AI145655 to X-P.K), VA Merit Review Award (I01BX003860 to C.E.H.) and VA Research Career Scientist Award (1IK6BX004607 to C.E.H). The funders had no role in study design, data collection and analysis, decision to publish, or preparation of the manuscript.

## Author Contributions

Conceptualization, C.U.; Supervision: C.U.; Investigation, C.U., R.F, L.C, K-W. C, K.L., W.Y., and J.Y.; Formal Analysis: C.U., A.N.; Resources: J.A, S.Z-P., H.Z., and X-P. K.; Writing – Original Draft, C.U.; Writing – Review & Editing, C.U., and C.E.H; Funding Acquisition, C.U and CEH.

## Supplementary Information

**S1 Fig. Sequence alignment of all Env SPs tested in this study**. Amino acid sequence alignment was performed using the Lasergene DNASTAR Megalign software. Within the alignment, dash (-) indicates missing residue.

**S2 Fig. Effect of SP swap on CMU06 infectivity, Env incorporation and lectin binding**. (A) Infectivity of 293T-derived CMU06 WT vs SP-swapped viruses in TZM.bl cells. (B) Measurement of Env incorporation by Western blots. (C) Binding of Env from CMU06 WT and SP-swapped viruses by gp120 mAbs and lectins (GNA and AAL) in Western blots. Densities of upper and lower Env bands (gray and black arrows) are shown.

**S3 Fig. CMU06 gp160 sequence**. N-glycosylation sites (red) are marked throughout the entire gp160 sequence that encompasses SP, constant regions (C1-C5), variable regions (V1-V5), and gp41.

**S4 Fig. Changes of neutralization sensitivity in the context of different Envs and SPs**. AUC changes of SP-swapped vs WT for the different virus strains tested are shown for comparison. NT, not tested.

**S5 Fig. Neutralization AUC values of WT and SP swapped viruses grown in 293T cells and PBMCs**. AUCs that decreased or increased by >30% and had p<0.05 relative to WT for the same MAbs or CD4-IgG are shown in orange or green, respectively. NT: not tested.

**S6 Fig. Glycan contents at glycosites on CMU06-WT (A), CMU06-398F1 (B), and CMU06-271**.**1 (C) produced in PBMCs vs 293T cells**. Glycosites detected on at least one of the Envs from PBMCs- and 293T-derived viruses are shown. (D) Percentages of unoccupied, complex, and oligomannose glycans on total detected glycosites (gp120 and gp41) from viruses grown in 293T cells vs PBMCs. The number of total glycosites for each virus is shown in parentheses.

**S7 Fig. SP prediction**. (A) SP prediction tool SignalP 3.0 applied to SPs of CMU06 WT and swap variants. Vertical red bars show the first amino acid after the cleavage site. (B) Probability of SP cleavage predicted by SignalP3.0. (C-D) PSIPRED-predicted MEMSAT-SVM helix orientation models (C) and DMPFold structures (D). One of the 5 structures predicted by PSIPRED for each SP is shown in panel D. α-helices: cyan, loops: magenta, residues C27, A29 and D31 around the cleavage site: magenta sticks.

