## Supplementary Information for "Signal Peptide of HIV-1 Envelope Modulates Glycosylation Impacting Exposure of V1V2 Epitopes"

**S1 Fig. Sequence alignment of all Env SPs tested in this study.** Amino acid sequence alignment was performed using the Lasergene DNASTAR Megalign software. Within the alignment, dash (-) indicates missing residue.

**S2 Fig. Effect of SP swap on CMU06 infectivity, Env incorporation and lectin binding**. (A) Infectivity of 293T-derived CMU06 WT vs SP-swapped viruses in TZM.bl cells. (B) Measurement of Env incorporation by Western blots. (C) Binding of Env from CMU06 WT and SP-swapped viruses by gp120 mAbs and lectins (GNA and AAL) in Western blots. Densities of upper and lower Env bands (gray and black arrows) are shown.

**S3 Fig. CMU06 gp160 sequence.** N-glycosylation sites (red) are marked throughout the entire gp160 sequence that encompasses SP, constant regions (C1-C5), variable regions (V1-V5), and gp41.

**S4 Fig. Changes of neutralization sensitivity in the context of different Envs and SPs.** AUC changes of SP-swapped vs WT for the different virus strains tested are shown for comparison. NT, not tested.

**S5 Fig. Neutralization AUC values of WT and SP swapped viruses grown in 293T cells and PBMCs.** AUCs that decreased or increased by >30% and had $p<0.05$ relative to WT for the same MAbs or CD4-IgG are shown in orange or green, respectively. NT: not tested.

**S6 Fig. Glycan contents at glycosites on CMU06-WT (A), CMU06-398F1 (B), and CMU06-271.1 (C) produced in PBMCs vs 293T cells.** Glycosites detected on at least one of the Envs from PBMCs- and 293T-derived viruses are shown. (D) Percentages of unoccupied, complex, and oligomannose glycans on total detected glycosites (gp120 and gp41) from viruses grown in 293T cells vs PBMCs. The number of total glycosites for each virus is shown in parentheses.

**S7 Fig. SP prediction**. (A) SP prediction tool SignalP 3.0 applied to SPs of CMU06 WT and swap variants. Vertical red bars show the first amino acid after the cleavage site. (B) Probability of SP cleavage predicted by SignalP3.0. (C-D) PSIPRED-predicted MEMSAT-SVM helix orientation models (C) and DMPFold structures (D). One of the 5 structures predicted by PSIPRED for each SP is shown in panel D. α-helices: cyan, loops: magenta, residues C27, A29 and D31 around the cleavage site: magenta sticks.

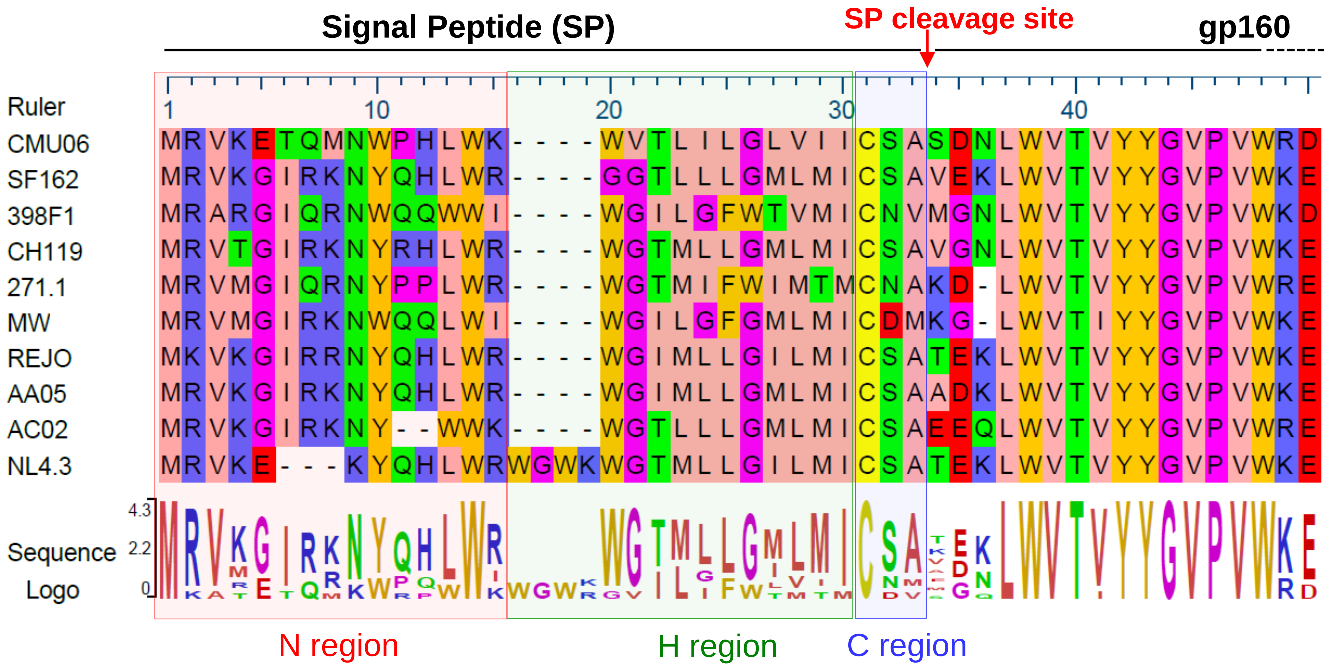


| A | B | C | D | E | F |
|---|---|---|---|---|---|
| CRF01_AE | 2 | Acute | 3 | 9 | 3 |
| B | 1A | Chronic | 6 | 7 | 4 |
| A | 1A | Acute | 3 | 8 | 1 |
| CRF07_BC | 2 | Chronic | 6 | 8 | 3 |
| CRF02_AG | 2 | Acute | 3 | 8 | 0 |
| C | 1A | 1A | 3 | 8 | 2 |
| B | 2 | T/F | 6 | 9 | 3 |
| B | - | Acute | 6 | 9 | 3 |
| B | - | Acute | 5 | 8 | 4 |
| B | 1 | Chronic | 5 | 10 | 3 |

Color Key:

| Physico-Chemical Property - Amino Acids |
|---|
| Aliphatic/hydrophobic - Alanine (A), Isoleucine (I), Leucine (L), Methionine (M), Valine (V) |
| Aromatic - Phenylalanine (F), Tryptophan (W), Tyrosine (Y) |
| Conformationally special - Glycine (G), Proline (P) |
| Cysteine (C) |
| Hydrophilic - Asparagine (N), Glutamine (Q), Serine (S), Threonine (T) |
| Negatively Charged - Aspartate (D), Glutamate (E) |
| Positively Charged - Arginine (R), Histidine (H), Lysine (K) |

A. Subtype
B. Neutralization tier; -, undetermined
C. Clinical infectious stage; T/F, transmitted/founder
D. # of positively charged residues in N region
E. # of hydrophobic residues in H region
F. # of leucine residues in H region

**S1 Fig**

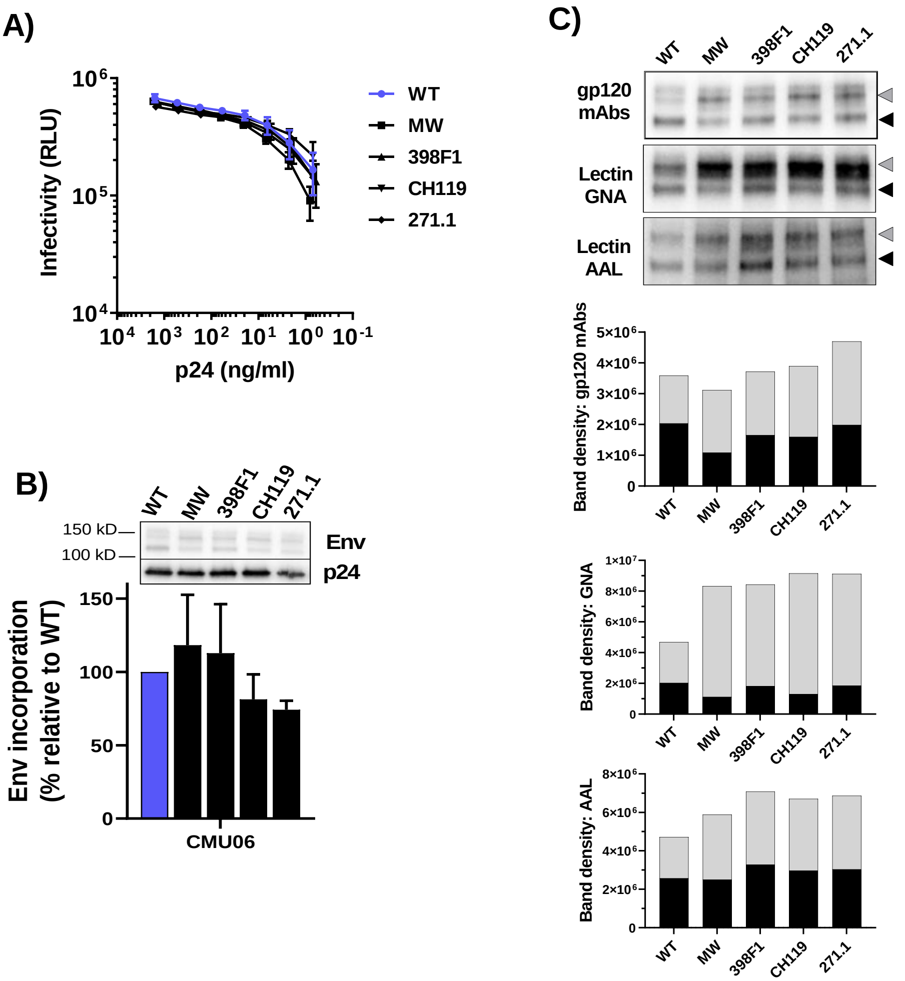
A)
Infectivity (RLU)
p24 (ng/ml)
WT
MW
398F1
CH119
271.1
B)
150 kD
100 kD
Env
p24
Env incorporation (% relative to WT)
CMU06
C)
gp120 mAbs
Lectin GNA
Lectin AAL
Band density: gp120 mAbs
Band density: GNA
Band density: AAL
Upper band
Lower band

S2 Fig

Signal peptide C1

MRVKETQMNWPHLWKWVTLILGLVIICSASDNLWVTVYYGVPVWRDADTTLFCASDAKAHETEVHNVW 70

ATHACVPTDPNPQEIHLE**N**VTENFNMWKNNMVEQMQEDVISLWDQSLKPCVKLTPLCVTLICINA**N**LTKI 140

V1 V2 C2

KSTTTDYNIG**N**ITDEVR**N**CSF**N**MTTEIIDKKQKVHALFYKLDIVPIENN**N**NSEYRLINC**N**TSVIKQACPKISF 215

DPIPIHYCAPAGYAILKCNDKNF**N**GTGPCK**N**VSSVQCTHGIKPVVSTQLLL**N**GSLAEEEIIIRSE**N**LTNNAKI 289

V3 C3

IIVHL**N**KSVEI**N**CTRPFNNVRTRMTMGPGHVFYRTGEITGDIRKAYCEI**N**GTKW**N**ETLKQVTEKLKEHFN 360

V4 C4

**N**KTIIFQPPSGGDLEITMHHFNCRGEFFYC**N**TTKLF**N**STWTE**N**GTMGGL**N**GTITLPCRIKQIVNMWQGA 429

V5 C5

GQAMYAPPINGKINCVS**N**ITGILLTRDGGV**N**NT**N**ETFRPGGGNIKDNWRSELYKYKVVQIEPLGIAPTRAK 501

gp41

RRVVEREKRAVGIGAMIFGFLGAAGSTMGAASITLTVQARQLLSGIVQQQSNLLRAIEAQQHMLQLTVW 572

GIKQLQARVLAVERYLKDQQFLGLWGCSGKIICTTAVPW**N**STWS**N**KSYEEIWD**N**MTWIEWEREIS**N**YTN 643

QIYEILTKSQDQQDRNEKDLLELDKWASLWNWF**N**ITNWLWYIKIFIMIVGGLIGLRIIFAVLSIVNRVRQGYS 716

PLSFQTLSHHQREPDRPEGIEEGGGEQGRDRSVRLVSGSRQKGDTKFILNA

**S3 Fig**

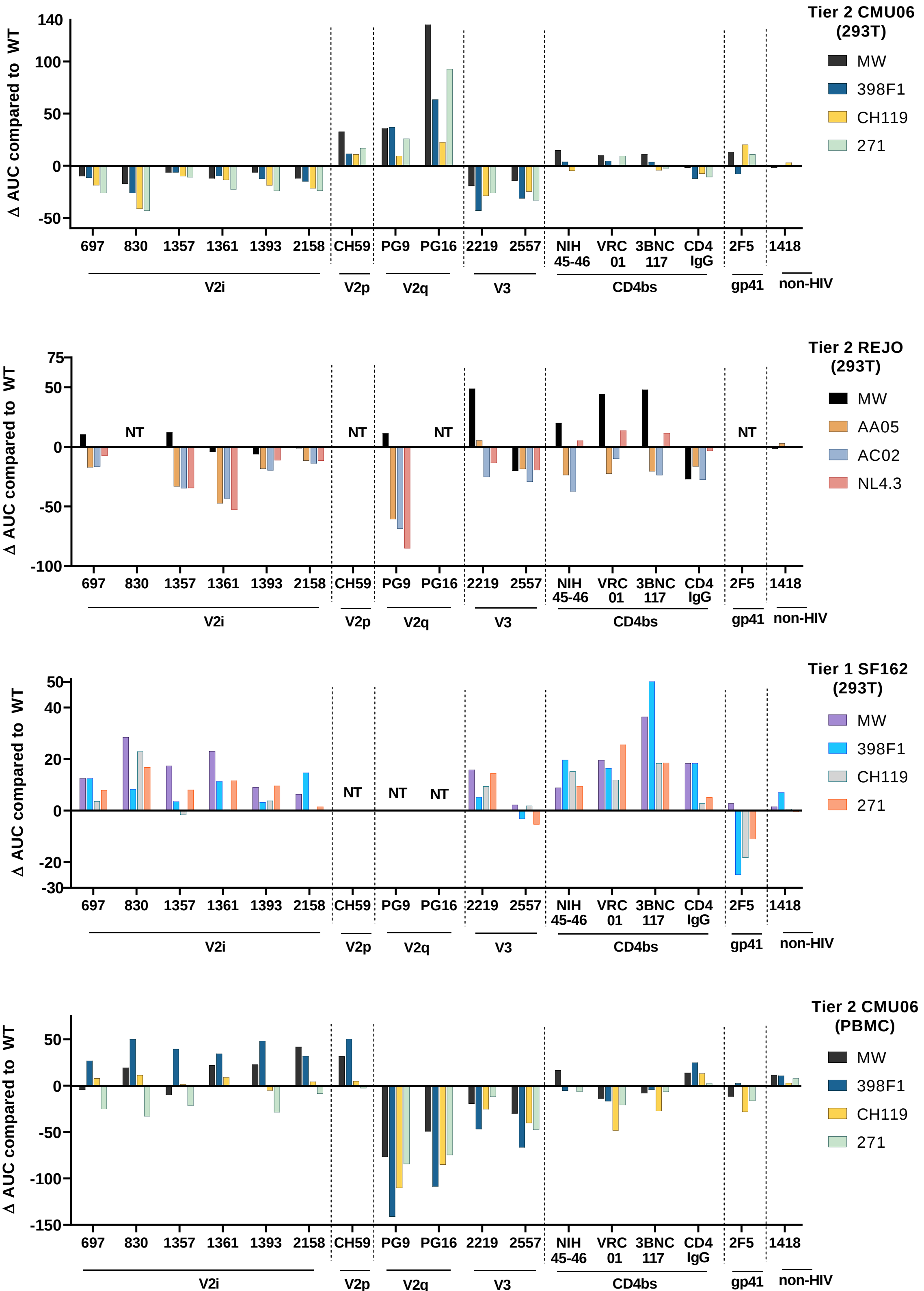
Tier 2 CMU06 (293T)
MW
398F1
CH119
271
Δ AUC compared to WT
140
100
50
0
-50
697
830
1357
1361
1393
2158
CH59
PG9
PG16
2219
2557
NIH 45-46
VRC 01
3BNC 117
CD4 IgG
2F5
1418
V2i
V2p
V2q
V3
CD4bs
gp41
non-HIV
Tier 2 REJO (293T)
MW
AA05
AC02
NL4.3
Δ AUC compared to WT
75
50
0
-50
-100
NT
Tier 1 SF162 (293T)
MW
398F1
CH119
271
Δ AUC compared to WT
50
40
20
0
-20
-30
NT
Tier 2 CMU06 (PBMC)
MW
398F1
CH119
271
Δ AUC compared to WT
50
0
-50
-100
-150

S4 Fig

| Env epitope | mAbs | CMU06 Neutralization: 293T (AUC) | | | | | REJO Neutraliation: 293T (AUC) | | | | | SF162 Viruses: 293T (AUC) | | | | | CMU06 Neutralization: PBMC (AUC) | | | | |
|---|---|---|---|---|---|---|---|---|---|---|---|---|---|---|---|---|---|---|---|---|---|
| | | WT | MW | 398F1 | CH119 | 271 | WT | MW | AA05 | AC02 | NL4.3 | WT | MW | 398F1 | CH119 | 271 | WT | MW | 398F1 | CH119 | 271 |
| V2i | 697 | 63 | 52 | 53 | 37 | 32 | 44 | 68 | 27 | 26 | 36 | 14 | 27 | 27 | 18 | 22 | 49 | 47 | 78 | 61 | 24 |
| | 830 | 140 | 132 | 102 | 85 | 88 | NT | NT | NT | NT | NT | 107 | 126 | 106 | 121 | 115 | 73 | 104 | 123 | 79 | 37 |
| | 1357 | 33 | 35 | 29 | 25 | 25 | 44 | 42 | 10 | 9 | 9 | 7 | 26 | 11 | 6 | 15 | 32 | 26 | 67 | 32 | 11 |
| | 1361 | 79 | 66 | 68 | 56 | 45 | 71 | 65 | 29 | 34 | 24 | 37 | 61 | 48 | 37 | 48 | 15 | 26 | 49 | 19 | 10 |
| | 1393 | 87 | 81 | 80 | 61 | 53 | 41 | 31 | 23 | 22 | 38 | 32 | 45 | 35 | 36 | 41 | 37 | 61 | 88 | 30 | 8 |
| | 2158 | 96 | 87 | 78 | 58 | 67 | 58 | 47 | 46 | 38 | 43 | 60 | 64 | 73 | 58 | 60 | 47 | 83 | 78 | 52 | 38 |
| V2p | CH59 | 162 | 194 | 173 | 173 | 179 | NT | NT | NT | NT | NT | NT | NT | NT | NT | NT | 83 | 117 | 141 | 100 | 105 |
| V2q | PG9 | 127 | 169 | 168 | 141 | 157 | 155 | 166 | 94 | 86 | 69 | NT | NT | NT | NT | NT | 168 | 91 | 26 | 57 | 83 |
| | PG16 | 15 | 132 | 69 | 40 | 94 | NT | NT | NT | NT | NT | NT | NT | NT | NT | NT | 123 | 73 | 14 | 37 | 48 |
| | PGT145 | NT | NT | NT | NT | NT | 354 | 333 | 317 | 320 | NT | NT | NT | NT | NT | NT | NT | NT | NT | NT | NT |
| V3 | 2219 | 47 | 28 | 4 | 18 | 21 | 82 | 125 | 82 | 51 | 63 | 171 | 186 | 176 | 180 | 185 | 51 | 32 | 5 | 26 | 40 |
| | 2557 | 61 | 47 | 30 | 36 | 29 | 89 | 66 | 71 | 64 | 68 | 175 | 177 | 172 | 177 | 170 | 78 | 48 | 12 | 38 | 31 |
| CD4bs | NIH45-46 | 206 | 215 | 178 | 186 | 194 | 134 | 154 | 110 | 97 | 139 | 146 | 159 | 170 | 166 | 160 | 81 | 98 | 70 | 81 | 74 |
| | VRC01 | 121 | 143 | 128 | 112 | 116 | 44 | 89 | 22 | 34 | 58 | 58 | 78 | 74 | 70 | 86 | 83 | 67 | 74 | 37 | 69 |
| | 3BNC117 | 166 | 186 | 176 | 167 | 185 | 151 | 199 | 131 | 127 | 163 | 107 | 144 | 157 | 126 | 126 | 119 | 114 | 108 | 90 | 106 |
| gp41 | 2F5 | 61 | 71 | 54 | 74 | 64 | NT | NT | NT | NT | NT | 38 | 47 | 19 | 26 | 33 | 48 | 36 | 52 | 20 | 31 |
| CD4 IgG | | 164 | 166 | 159 | 153 | 155 | 118 | 78 | 103 | 86 | 110 | 166 | 185 | 185 | 169 | 171 | 102 | 119 | 130 | 115 | 107 |
| non-HIV | 1418 | 9 | 3 | 9 | 6 | 8 | 10 | 8 | 12 | 10 | 9 | 3 | 4 | 4 | 3 | 2 | 1 | 12 | 21 | 4 | 10 |

Decrease by >30%, p<0.05

Increase by >30%, p<0.05

**S5 Fig**

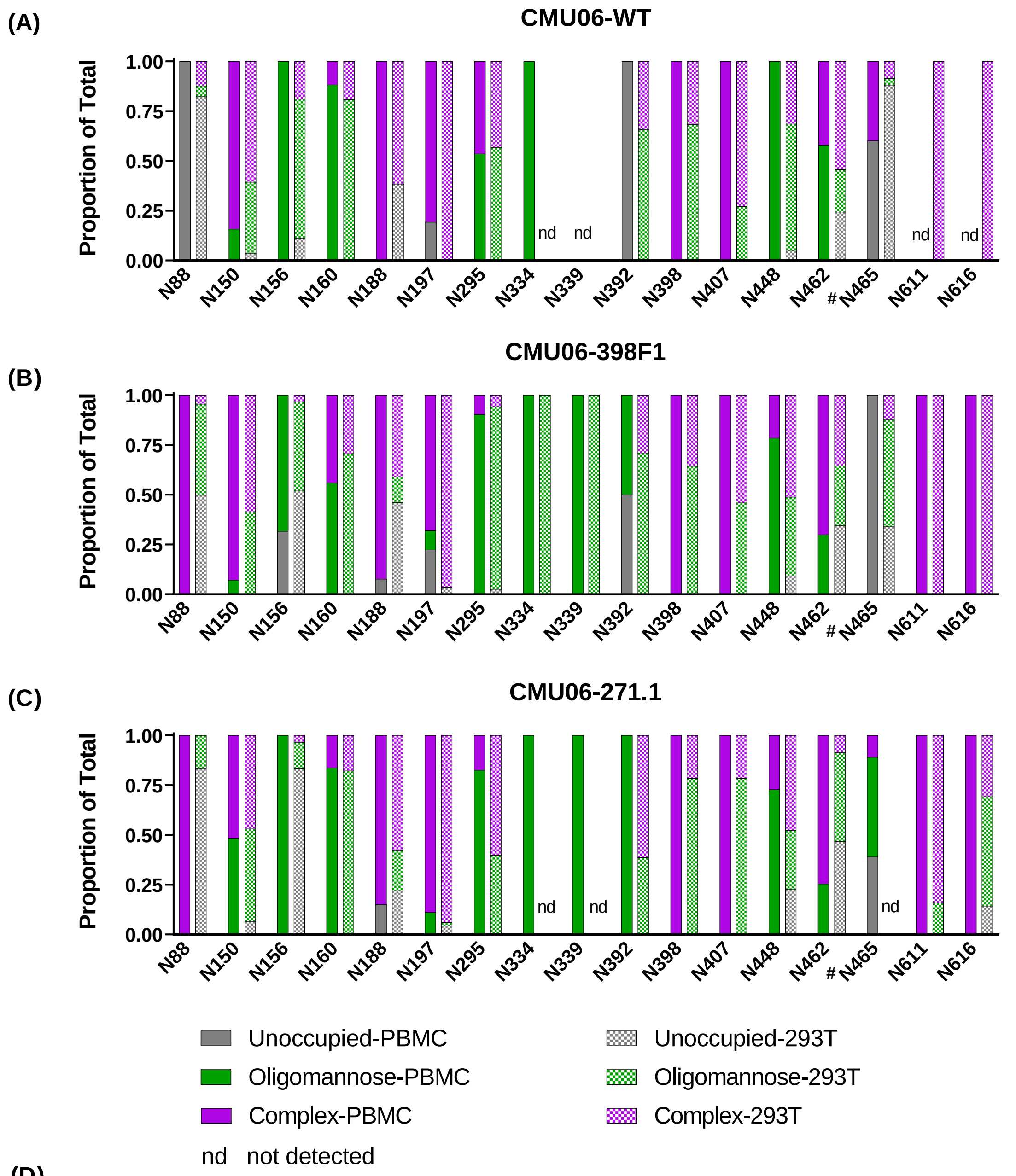


(D)

| Host cells | % Unoccupied (total PNGS) | | | % Complex (total PNGS) | | | % Oligomannose (total PNGS) | | |
|---|---|---|---|---|---|---|---|---|---|
| | WT | 398F1 | 271.1 | WT | 398F1 | 271.1 | WT | 398F1 | 271.1 |
| **293T** | 16.4 (16) | 15.1 (16) | 19.1 (16) | 48.2 (16) | 45.1(16) | 37.6 (16) | 35.4 (16) | 39.8 (16) | 43.3 (16) |
| **PBMC** | 23.7 (16) | 17.3 (18) | 5.5 (19) | 37.8 (16) | 49.9 (18) | 45.9 (19) | 38.5 (16) | 32.8 (18) | 48.6 (19) |

**S6 Fig**

A

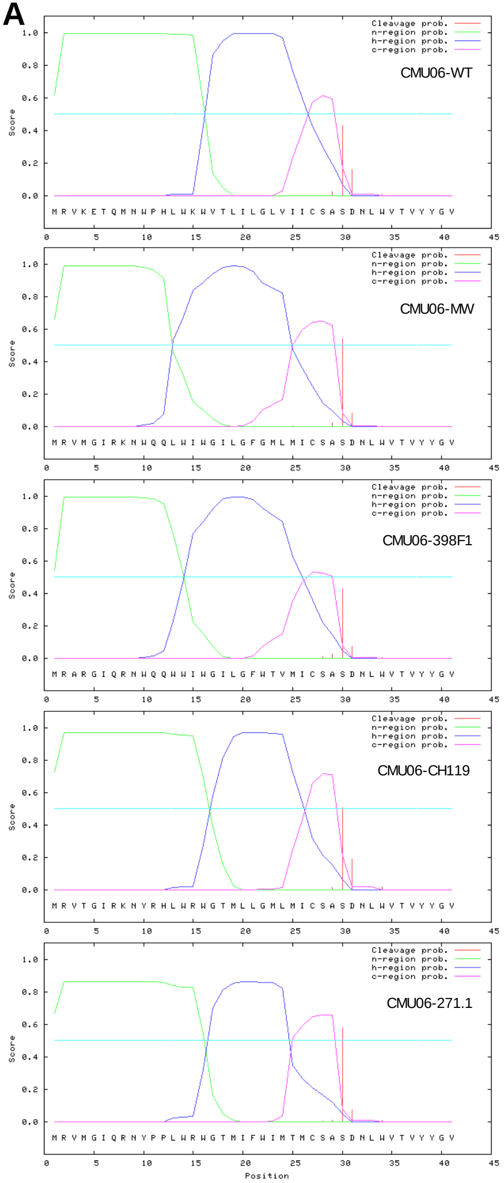


B

| Probability | CMU06-WT | CMU06-MW | CMU06-398F1 | CMU06-CH119 | CMU06-271.1 |
|---|---|---|---|---|---|
| Max Cleavage site | 0.427 | 0.542 | 0.428 | 0.504 | 0.577 |
| Signal peptide | 0.62 | 0.655 | 0.538 | 0.727 | 0.668 |
| Signal anchor | 0.375 | 0.333 | 0.457 | 0.243 | 0.192 |

C

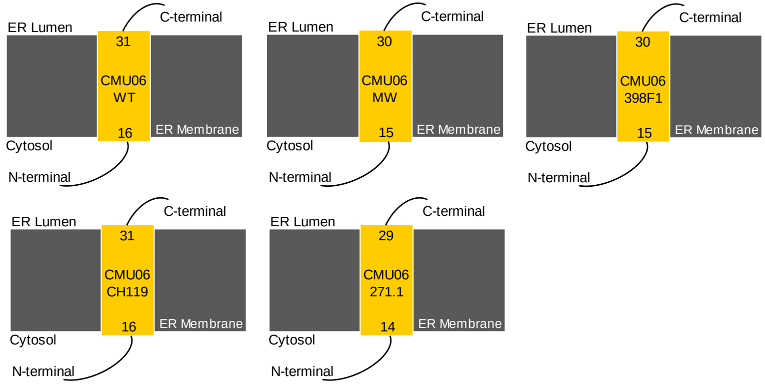


D

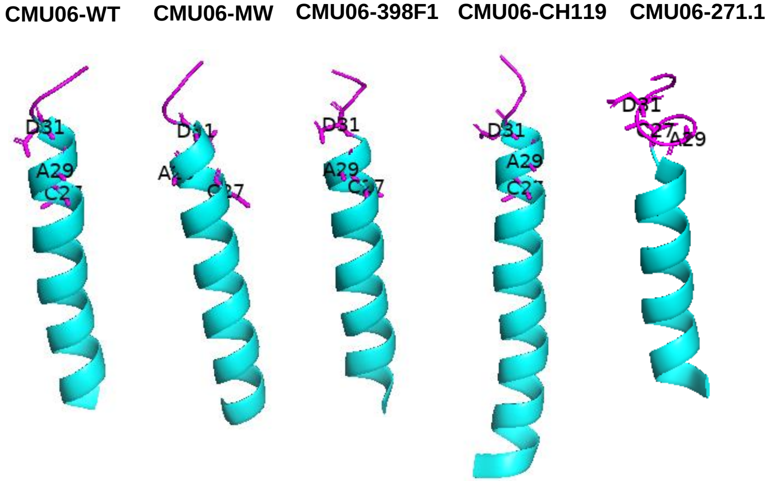


**S7 Fig**
